# Tissue-specific transcriptome profiling of the *Arabidopsis thaliana* inflorescence stem reveals local cellular signatures

**DOI:** 10.1101/2020.02.10.941492

**Authors:** Dongbo Shi, Virginie Jouannet, Javier Agustí, Verena Kaul, Victor Levitsky, Pablo Sanchez, Victoria V Mironova, Thomas Greb

**Affiliations:** Department of Developmental Physiology, Centre for Organismal Studies (COS), Heidelberg University, Im Neuenheimer Feld 230, 69120 Heidelberg, Germany; Gregor Mendel Institute (GMI), Austrian Academy of Sciences, Vienna Biocenter (VBC), Dr. Bohr-Gasse 3, 1030 Vienna, Austria; Instituto de Biología Molecular y Celular de Plantas (IBMCP), l’Enginyer, Carrer de l’Enginyer Fausto Elio, 24, 46011. Universitat Politècnica de València. València, Spain; Novosibirsk State University, Novosibirsk 630090, Russia; Institute of Cytology and Genetics SB RAS, Novosibirsk 630090, Russia

## Abstract

Genome-wide gene expression maps with a high spatial resolution have substantially accelerated molecular plant science. However, the number of characterized tissues and growth stages is still small because of the limited accessibility of most tissues for protoplast isolation. Here, we provide gene expression profiles of the mature inflorescence stem of *Arabidopsis thaliana* covering a comprehensive set of distinct tissues. By combining fluorescence-activated nucleus sorting and laser-capture microdissection with next generation RNA sequencing, we characterize transcriptomes of xylem vessels, fibers, the proximal and the distal cambium, phloem, phloem cap, pith, starch sheath, and epidermis cells. Our analyses classify more than 15,000 genes as being differentially expressed among different stem tissues and reveal known and novel tissue-specific cellular signatures. By determining transcription factor binding regions enriched in promoter regions of differentially expressed genes, we furthermore provide candidates for tissue-specific transcriptional regulators. Our datasets predict expression profiles of an exceptional amount of genes and allow generating hypotheses toward the spatial organization of physiological processes. Moreover, we demonstrate that information on gene expression in a broad range of mature plant tissues can be established with high spatial resolution by nuclear mRNA profiling.

**One sentence summary:** A genome-wide high-resolution gene expression map of the Arabidopsis inflorescence stem is established.

## Introduction

Characterizing gene expression in individual cell types is a powerful tool for revealing local molecular signatures in multicellular organisms. By combining genetically encoded fluorescent reporters driven by tissue-specific promoters and fluorescence-activated cell sorting (FACS), high-resolution gene expression profiles have been established for developmental hotspots like the root and shoot tips of *Arabidopsis thaliana* (Arabidopsis) (Birnbaum et al., 2003; Brady et al., 2007; Yadav et al., 2009; Yadav et al., 2014). Although these datasets have served as central resources for the scientific community for many years, high-resolution gene expression maps have not been developed for many other organs or tissues. One of the obstacles is the reliable isolation of RNA from more differentiated tissues and cell types. Depending on their identity or developmental stage, plant cells are surrounded by cell walls with very diverse properties. This requires extensive tuning of methods for disrupting cell walls for each case individually (Bart et al., 2006; Lin et al., 2014) or hampers the isolation overall. Even when protoplasts can be isolated, their highly diverse sizes render the subsequent sorting process challenging. Laser capture microdissection (LCM) is an alternative tool for a precise isolation of cell material (Schad et al., 2005; Chandran et al., 2010; Agusti et al., 2011; Blokhina et al., 2016). Spatial specificity of the profiling process is lower in this case, because genetically encoded markers for cell identity can hardly be used. However, LCM is a powerful method when genetically encoded markers are not available and cell types can be clearly identified by morphology or anatomical position.

Plant stems play fundamental roles in determining shoot architecture and act as transport routes connecting photosynthetically active source organs with the remaining plant body (Sanchez et al., 2012; Serrano-Mislata and Sablowski, 2018). After the transition from vegetative to reproductive growth, Arabidopsis forms inflorescence stems, which are composed of various tissues including epidermis, cortex, starch sheath, vascular bundles and pith (Figures. 1A, B). These tissues fulfil very special roles in plant bodies and, by acting in concert, form the stem as a functional unit. For example, the epidermis protects the plant from desiccation by building a transpiration barrier and serves as a first line of defense against pathogens (Esau, 1977; Suh et al., 2005). In turn, the starch sheath, also designated as endodermis, executes gravity sensing (Morita et al., 2002; Nawrath et al., 2013). In vascular bundles, xylem and phloem tissues are composed of specialized cell types such as xylem fibers, xylem vessel elements, phloem sieve elements and phloem companion cells enabling water and nutrient transport (Figure. 1B). In comparison to these highly specialized cells, cambium stem cells maintain the potential to generate secondary xylem and phloem cells for increasing transport capacities and mechanical support of the growing shoot system (Suer et al., 2011). Interestingly, many of the tissues found in stems are also found in other organs like roots or leaves. However, their organ-specific profiles have not been characterized systematically.

**Figure 1.**
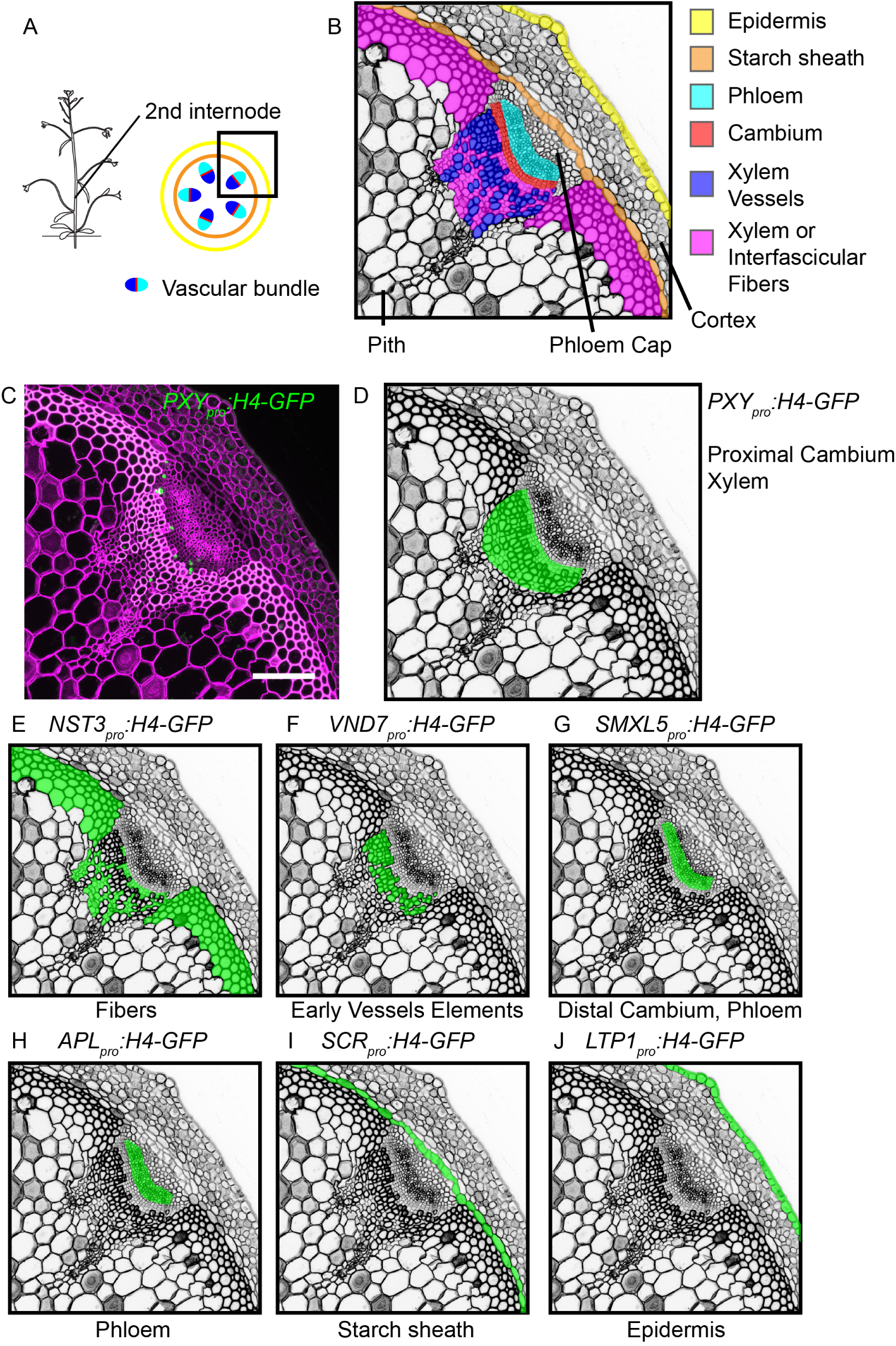
Expression patterns of *H4-GFP* transgenes in the second internode. (A, B) Representations of tissue configuration in the second bottom-most Arabidopsis internode as seen in cross sections. The region shown in B corresponds to the black squared region in A. (C) Maximum intensity projection of confocal images of internode sections of the *PXY_pro_:H4-GFP* line. The GFP signal is shown in green and cell walls are stained by Direct Red 23 and visualized in magenta. Scale bar: 100 µm. Note that only nuclei in the limited observable depth of the section are visualized. (D-J) Schematic indication of activity patterns of the different *H4-GFP* transgenes in cross sections of the second internode. Original data are shown in Supplemental Figure 1.

Like in most dicotyledonous species, Arabidopsis stems undergo a major anatomical transition during the initiation of radial growth and of extensive wood formation (Sanchez et al., 2012). In stems holding a primary tissue conformation, cambium stem cells are restricted to vascular bundles whereas in secondary stems, cambium cells are also found in interfascicular regions. Thereby, cambium stem cells establish concentric domains of vascular tissues important for an organized radial growth (Sehr et al., 2010; Sanchez et al., 2012).

Considering the central role of plant stems in determining plant architecture and physiology, it is vital to have information on gene expression profiles from a comprehensive set of cell types and tissues. From these data, physiological and developmental features of stem tissues can be reveled and the organ can be characterized as a functional unit. Although several studies have provided transcriptome profiles from several stem tissues and stages (Ko and Han, 2004; Ko et al., 2004; Suh et al., 2005; Brackmann et al., 2018), a systematic analysis has not been performed and stem-specific tissues like the pith or the phloem cap have not been targeted at all.

Due to the large organ diameter and the heterogeneity of cell walls in plant stems, enzymatic protoplasting seems unsuitable for harvesting material from different stem tissues with the same efficiency. Therefore, nucleus isolation intrudes itself as an attractive alternative. Nuclei are released from tissues after manual chopping (Galbraith et al., 1983) and are possible to be isolated from individual tissues by fluorescence-activated nucleus sorting (FANS) (Zhang et al., 2005; Zhang et al., 2008; Slane et al., 2014). Moreover, although nuclear mRNA and cytosolic mRNA have different compositions and roles (Yang et al., 2017; Choudury et al., 2019), previous studies demonstrated that cellular gene expression can be deduced accurately from nuclear mRNA levels (Zhang et al., 2008; Deal and Henikoff, 2010; You et al., 2017).

Motivated by these considerations, we employed in this study FANS and LCM to extract and profile mRNA from a large set of tissues present in the primary Arabidopsis stem. By using tissue-specific promoters for fluorescence labeling and profiling nuclei from seven tissues and LCM for two tissues for which no promoter has been identified, we reveal spatial information on gene activities in a genome-wide fashion. Because the primary inflorescence stem contains a large spectrum of tissues including extremes like cambium stem cells and terminally differentiated cells in the vasculature, our results demonstrate the broad applicability of these approaches.

## Results

### Establishment of plant lines for tissue-specific labelling of nuclei

To establish experimental access to mRNA from individual stem tissues, we first screened literature for promoters specifically active in distinct tissues (Schürholz et al., 2018). As a result, the *NAC SECONDARY WALL THICKENING PROMOTING3* (*NST3*) promoter was chosen to label fibers (Mitsuda et al., 2007), the *VASCULAR RELATED NAC DOMAIN PROTEIN7* (*VND7*) promoter for differentiating vessel elements (Kubo et al., 2005; Yamaguchi et al., 2010), the *PHLOEM INTERCALATED WITH XYLEM* (*PXY*)*/TDIF RECEPTOR* (*TDR*) promoter for differentiating xylem cells and proximal cambium cells (Fisher and Turner, 2007; Hirakawa et al., 2008; Shi et al., 2019), the *SMAX1-LIKE5* (*SMXL5*) promoter for differentiating phloem cells and distal cambium cells (Wallner et al., 2017; Shi et al., 2019), the *ALTERED PHLOEM DEVELOPMENT (APL)* promoter for differentiated phloem cells (Bonke et al., 2003), the *SCARECROW* (*SCR*) promoter for starch sheath cells (Wysocka-Diller et al., 2000), and the *LIPID TRANSFER PROTEIN1* (*LTP1*) promoter for epidermis cells (Baroux et al., 2001) (Figure 1B). To confirm tissue-specificity of promoter activities, we generated transgenic plant lines expressing a fusion protein between histone H4 and the Green Fluorescent Protein (H4-GFP) under the control of each promoter (*GENE_pro_:H4-GFP*), respectively. Microscopic inspection of cross sections from the second bottom-most internode showed that only nuclei in the expected tissues were labeled by GFP (Figures 1C-J, Supplemental Figure 1) and, thus, that our lines carried GFP-positive nuclei in a tissue-specific manner.

### Tissue-specific gene activity in stems can be determined by nucleus profiling

To see whether we were able to faithfully extract tissue-specific mRNA, we first focused on inner tissues and tissues producing prominent secondary cell walls expecting that isolation was most challenging in these cases. Therefore, we collected nuclei from fibers marked by *NST3_pro_* activity, the distal cambium (*SMXL5_pro_*) and the phloem (*APL_pro_*) by manually chopping the second bottom-most internode of inflorescence stems and subsequent FANS. From each line, GFP-positive and negative nuclei were harvested (Figures 2A-C, Supplemental Figures 2E-I) and 15,000 nuclei per sample were processed for transcriptome analyses (Figures 2A-C). Our procedure included, SMART-seq2 amplification of mRNA (Picelli et al., 2013) and RNA-seq analysis of three replicates for each sample type. Following this strategy, we found that the levels of the *H4-GFP* mRNA (detected through the *GFP* sequence) and the mRNA of the *NST3, SMXL5* or *APL* genes whose promoters were used for expressing H4-GFP in the respective lines, were significantly enriched in extracts from GFP-positive nuclei compared to extracts from GFP-negative nuclei in all cases (Figures 2D-I, Supplemental Dataset 1). These findings indicated that cell type-specific transcriptomes were thoroughly accessible following our experimental pipeline.

**Figure 2.**
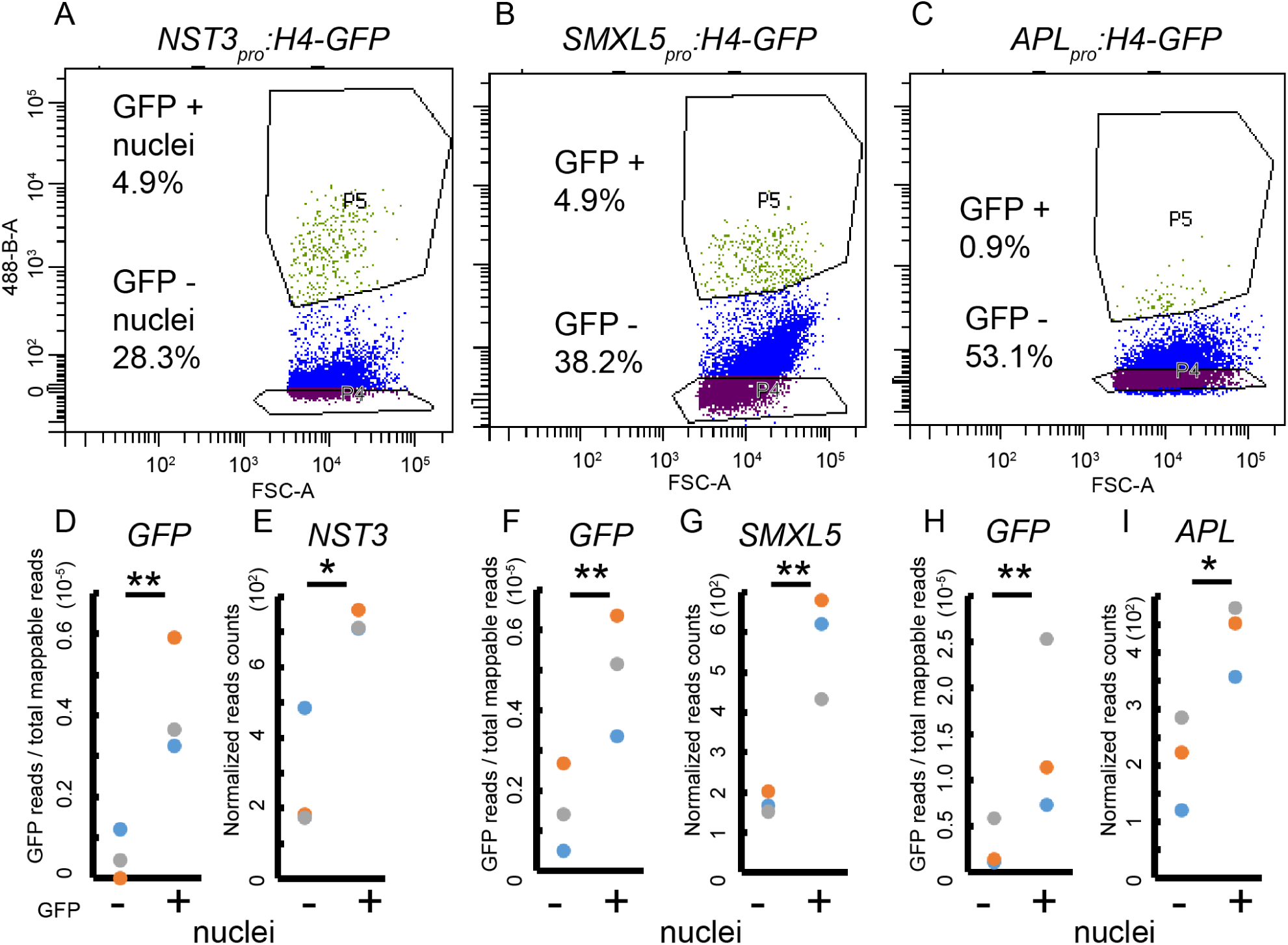
Sorting gates and gene expression analyses of GFP-positive or GFP-negative nuclei. (A-C) Plot of the gate settings defining GFP-positive nuclei (P5) and GFP-negative nuclei (P4) while sorting nuclei from the second internode of *NST3_pro_:H4-GFP* (A), *SMXL5_pro_:H4-GFP* (B) and *APL_pro_:H4-GFP* (C) lines, respectively. The ratios of each population compared to all particle counts are labeled. The X axis (FSC; forward scatter intensity) indicates the diameter of nuclei, the Y axis (488) indicates GFP fluorescence. (D, F, H) Comparison of read counts mapped to the *GFP* sequence to the total number of mappable reads to the *Arabidopsis* genome in GFP-positive and GFP-negative nuclei for each transgenic line. n = 3 for each population for each line. ** indicates *p* < 0.01 in the inverted beta-binomial test. (E, G, I) Normalized read counts for *NST3*, *SMXL5*, and *APL* genes in GFP-positive and GFP-negative nuclei for each transgenic line, respectively. *n* = 3 for each population. ** indicates *p* < 0.01 and * indicates *p* < 0.05 determined by the Wald test.

### Transcriptome analysis of seven stem tissues

Next, we performed transcriptome profiling of GFP-positive nuclei harvested from the remaining four lines (*PXY_pro_*, *LTP1_pro_*, *VND7_pro_*, *SCR_pro_*, Supplemental Figure 2A-D) by RNA-seq and contrasted obtained transcriptome data from GFP-positive nuclei of all seven tissues (Supplemental Figure 3, Supplemental Dataset 2). After having classified 4 out of 21 datasets as outliers based on principle component (PCA) and correlation analyses (Supplemental Figure 3), we kept 17 datasets from which replicates clustered and correlated as expected (n=2 or 3, Figures 3A, B, Supplemental Dataset 2). Confirming the biological relevance of the obtained profiles, xylem-related (*PXY_pro_*, *NST3_pro_*, *VND7_pro_*) and phloem-related (*SMXL5_pro_*, *APL_pro_*) datasets showed a high correlation coefficient among each other but belonged to two different major branches within the correlation plot (Figure 3B).

**Figure 3.**
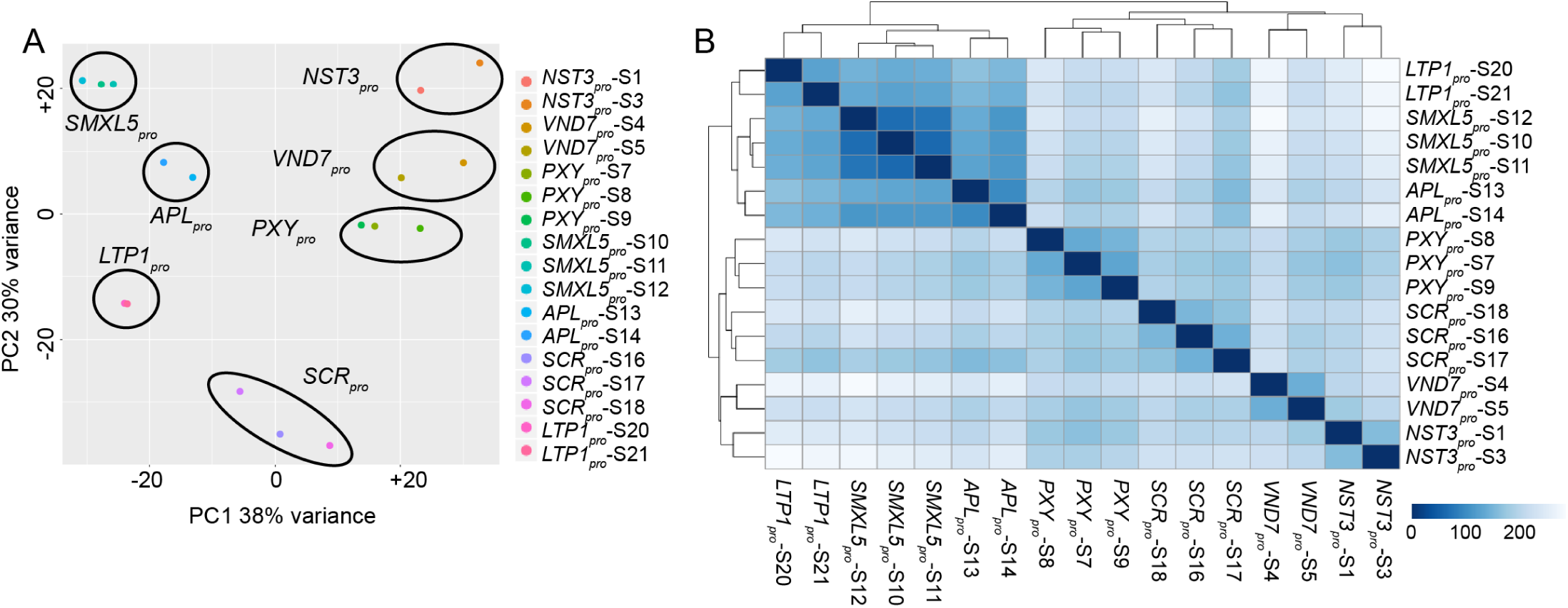
Statistical comparisons of RNA-seq datasets derived from GFP-positive nuclei of seven stem tissues. (A) PCA analysis on log-transformed normalized read counts of each RNA-seq dataset. (B) Heatmap displaying the statistical distance between each RNA-seq dataset according to the color code. Two or three replicates were obtained from each nucleus population.

Overall, among 37,051 Arabidopsis protein-coding and non-coding genes (Cheng et al., 2017), we detected 25,679 – 33,949 genes to be expressed in the respective tissues (Transcripts Per Million, TPM > 1), suggesting sufficient coverage by our RNA-seq analyses (Table 1). The average percentage of 14.4 % to 24.3 % of reads mapping to introns compared to reads mapping to exons was higher than the 7.7 % found on average in our previous whole RNA-seq analyses of the second internode (Brackmann et al., 2018). This difference suggested that nucleus-derived RNA contains more non-processed transcripts compared to RNA derived from whole tissues (Table 1). In contrast, the average ratio of reads mapping to intergenic regions compared to the ratio of reads mapping to exons was comparable (Table 1) demonstrating that the observed difference in the percentage of intron-associated reads did not result from contaminations by genomic DNA.

**Table 1.**
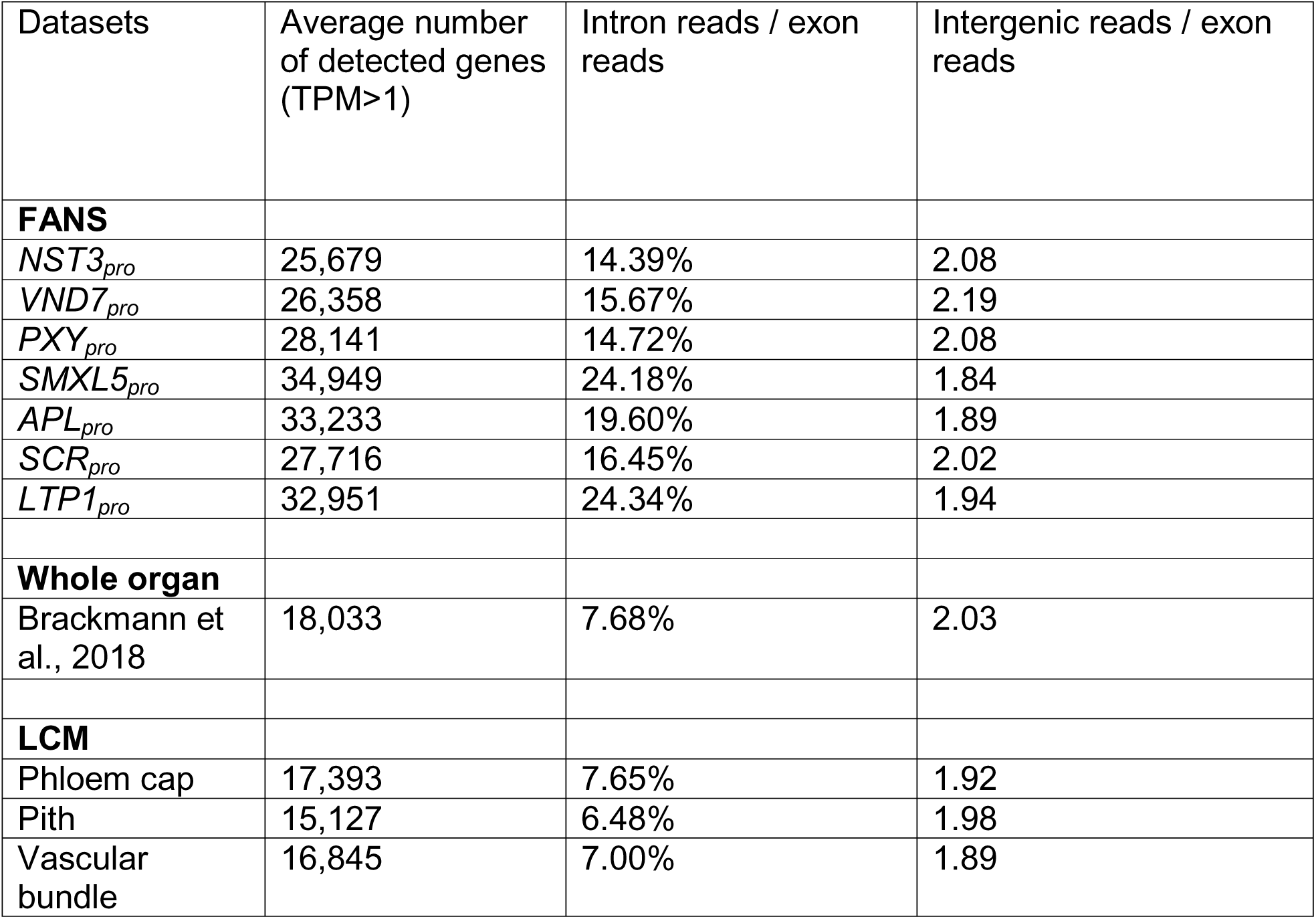
Summary of RNA-seq results. Summary of RNA-seq results for the different tissues from the second internode collected by FANS, for the whole organ by conventional RNA extraction (Brackmann et al., 2018) and collected by LCM. n = 2 or 3 for each dataset.

| Datasets | Average number of detected genes (TPM>1) | Intron reads / exon reads | Intergenic reads / exon reads |
| --- | --- | --- | --- |
| **FANS** |  |  |  |
| *NST3<sub>pro</sub>* | 25,679 | 14.39% | 2.08 |
| *VND7<sub>pro</sub>* | 26,358 | 15.67% | 2.19 |
| *PXY<sub>pro</sub>* | 28,141 | 14.72% | 2.08 |
| *SMXL5<sub>pro</sub>* | 34,949 | 24.18% | 1.84 |
| *APL<sub>pro</sub>* | 33,233 | 19.60% | 1.89 |
| *SCR<sub>pro</sub>* | 27,716 | 16.45% | 2.02 |
| *LTP1<sub>pro</sub>* | 32,951 | 24.34% | 1.94 |
| **Whole organ** |  |  |  |
| Brackmann et al., 2018 | 18,033 | 7.68% | 2.03 |
| **LCM** |  |  |  |
| Phloem cap | 17,393 | 7.65% | 1.92 |
| Pith | 15,127 | 6.48% | 1.98 |
| Vascular bundle | 16,845 | 7.00% | 1.89 |

When comparing expression levels in the different tissues, reads from the *APL, SCR, NST3, VND7, PXY* and *SMXL5* genes were, as expected, enriched in samples derived from GFP-positive nuclei from the respective *GENE_pro_:H4-GFP* lines (Figures 4A, D, F, H, K, M). Moreover, *APL* peaked together with the *SIEVE-ELEMENT-OCCLUSION-RELATED1 (SEOR1)* and *NAC DOMAIN-CONTAINING PROTEIN86 (NAC086)* genes, which are known to be expressed in phloem cells (Froelich et al., 2011; Furuta et al., 2014) (Figures 4B, C). Reads of the starch sheath-expressed *PIN-FORMED3 (PIN3)* gene (Friml et al., 2002), were also most abundant in *SCR_pro_*-positive nuclei (Figure 4E). Likewise, reads from *NST1*, known to be expressed in fiber cells (Mitsuda et al., 2005) had a maximum in *NST3_pro_*-positive nuclei (Figure 4G) and reads from the *VND6* gene, the closest homologue of *VND7* (Zhong et al., 2008) and its downstream target *XYLEM CYSTEINE PROTEASE1 (XCP1)* (Zhong et al., 2010; Yamaguchi et al., 2011) showed the highest activity in *VND7_pro_*-positive nuclei (Figures 4I, J). We also found activity of *WUSCHEL RELATED HOMEOBOX4* (*WOX4*) (Figure 4L), whose expression domain is congruent with the *PXY* expression domain (Hirakawa et al., 2008; Suer et al., 2011; Brackmann et al., 2018; Shi et al., 2019) to peak in *PXY_pro_*-positive nuclei. In addition, *MORE LATERAL GROWTH1* (*MOL1*)*, PHLOEM EARLY DOF1* (*PEAR1*) and *PEAR2* are expressed in *SMXL5_pro_*-positive cells (Gursanscky et al., 2016; Miyashima et al., 2019) and peaked, as expected, in *SMXL5_pro_*-positive nuclei (Figures 4N-P). Although *LTP1* reads were not enriched in *LTP1_pro_*-positive nuclei (Figure 4Q), the reads of *FIDDLEHEAD (FDH)* and *ECERIFERUM6 (CER6)*, which are known to be expressed in the epidermis (Yephremov et al., 1999; Pruitt et al., 2000; Hooker et al., 2002), were most abundant in *LTP1_pro_*-positive nuclei (Figures 4R, S). Further verifying that we succeeded in determining the transcriptome of the epidermis, we found that 32 out of 40 genes identified previously to show the highest activity levels in the stem epidermis (Suh et al., 2005) had more normalized read counts in the *LTP1_pro_*-derived dataset compared to the overall tissue-average and the reads of 23 out of the same 40 genes were indeed most abundant in *LTP1_pro_*_-_positive nuclei (Supplemental Figure 4).

**Figure 4.**
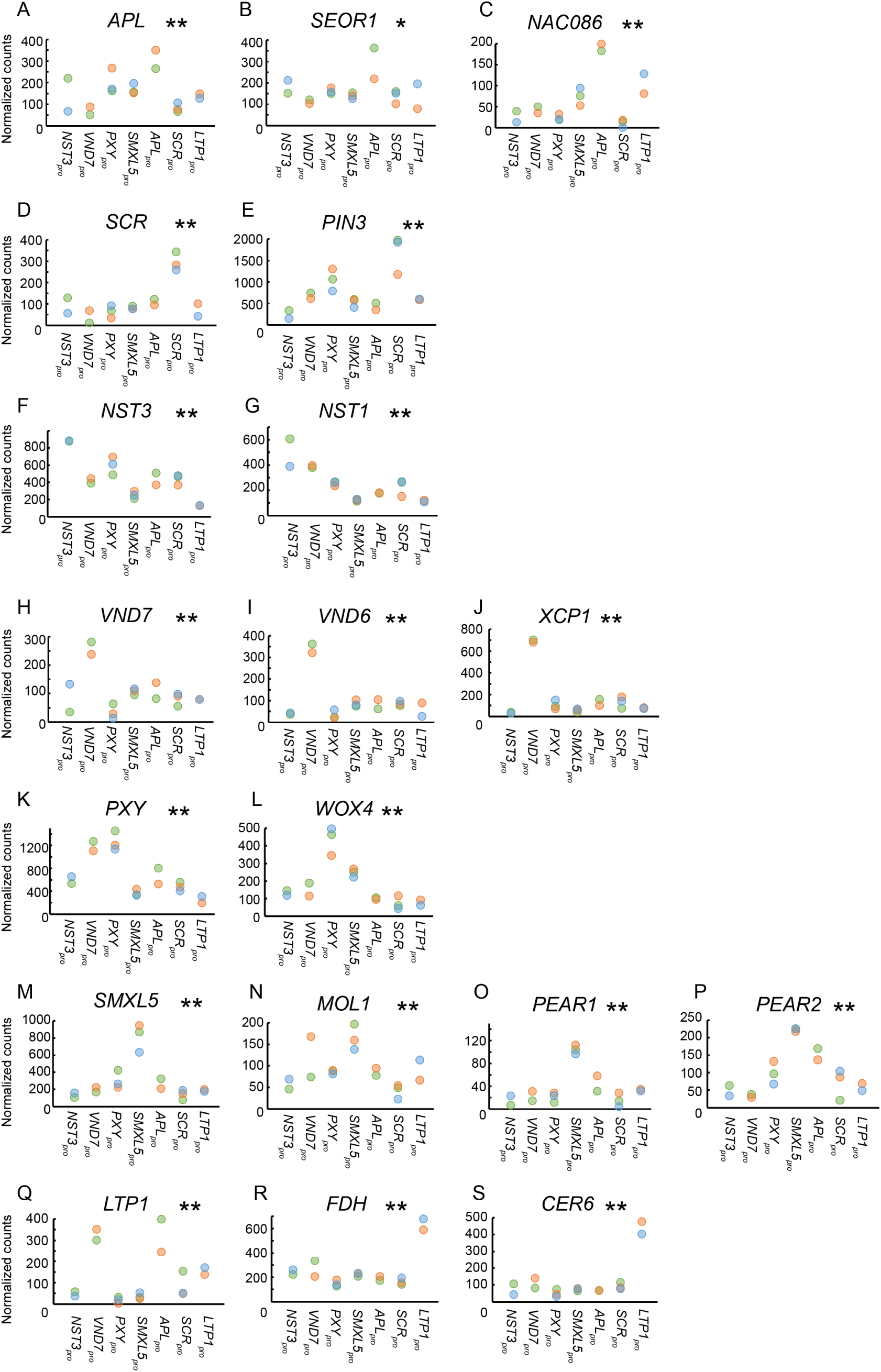
Transcriptome dataset comparisons of seven stem tissues. (A-S) Normalized gene read counts of the indicated genes among seven different tissues displayed for each replicate individually. *, ** indicates p < 0.05 and 0.01, respectively in LRT. Please note that the null hypothesis to be rejected in the LRT is that genes have similar expression among the seven different tissues.

### Transcriptome dataset of the phloem cap and the pith

Because for the pith and the phloem cap no reliable tissue-specific promoters have been identified to date, we employed LCM to determine transcriptome profiles of these tissues. To this end, we first collected the phloem cap and the pith followed by the collection of the remaining vascular bundle which we harvested as a comparison (Figure 5A, n = 2 replicates). Subsequently, RNA was extracted, mRNA was amplified and analyzed by RNA-seq (Figure 5A, Supplemental Dataset 3). Confirming reliable sample preparation, replicates generated from each sample type grouped together in PCA plots (Figure 5B). Moreover, correlation analyses showed that the phloem cap profile and the profile from the remaining vascular bundle were more similar to each other than to the pith profile (Figure 5C). This finding was expected considering the high spatial and ontogenetic relatedness between the vascular bundle and the phloem cap.

**Figure 5.**
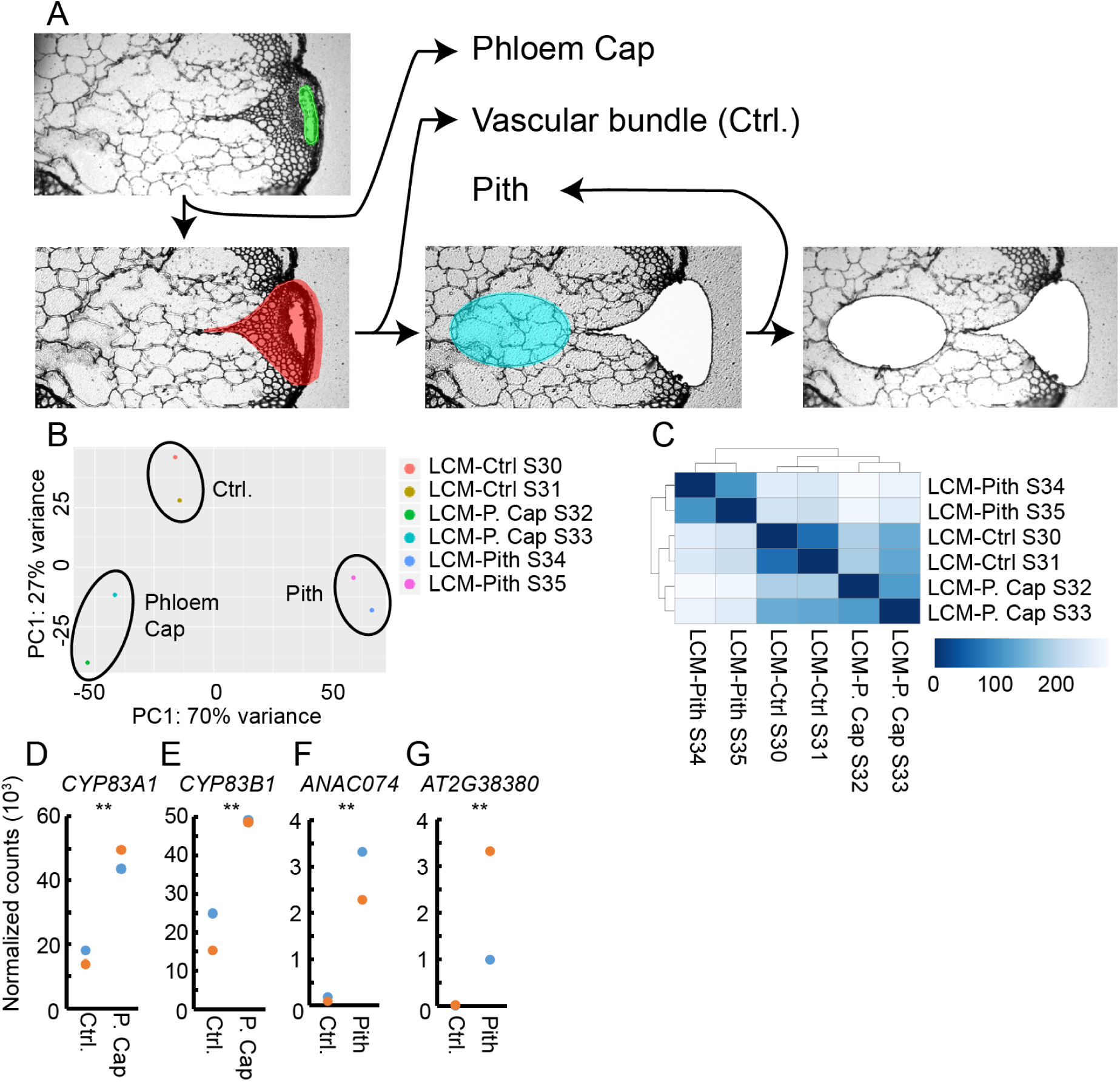
Transcriptome analysis of the phloem cap and the pith using LCM. (A) Strategy of tissue collection using LCM shown for one individual sample. (B) PCA analysis on log-transformed normalized read counts of each LCM-derived RNA-seq dataset. (C) Heatmap displaying the statistical distance between each RNA-seq dataset. Two replicates were obtained from each tissue type. (D-G) Normalized gene read counts in comparing phloem cap / pith and the vascular bundle (Ctrl.). ** indicates *p* < 0.01 in Wald tests.

On average, we detected 15,127 – 17,393 genes to be expressed in each LCM-derived sample type (TPM > 1) suggesting a lower coverage in comparison to our FANS-based analyses (Table 1). The average ratio of the number of reads mapped to intron regions compared to the number of reads mapped to exons for each sample type were with 6.5 % to 7.7 % in a similar range as in standard RNA-seq approaches for the Arabidopsis stem (Brackmann et al., 2018) (Table 1). Supporting reliable profiling, reads from the *CYTOCHROME P450 83A1 (CYP83A1)* and *CYP83B1* genes, which are part of the indole and aliphatic glucosinolate biosynthetic pathways and known to be expressed in the phloem cap (Nintemann et al., 2018) were significantly enriched in phloem cap samples (Figures 5D, E). In addition, reads from the *ANAC074* and *AT2G38380* genes which are known to be expressed in the pith (Fujimoto et al., 2018; Schürholz et al., 2018) were enriched in pith-derived samples (Figures 5F, G).

### Identifying genes with tissue-related expression patterns

To identify genes predominantly active in individual tissues, we next applied the DESeq2 software package and the likelihood ratio test (LRT) to our FANS-derived datasets (Love et al., 2014). Thereby, we classified 14,063 genes as significantly differentially expressed (SDE) genes (Supplemental Datasets 4, 5; Benjamini-Hochberg (BH) adjustment of *p* value in LRT < 0.01). Based on their activity profiles, SDE genes were categorized into 93 clusters using hierarchical clustering (Figure 6, Supplemental Figure 5, Supplemental Dataset 6). Clusters contained 9 to 1,271 genes with 151 genes on average which, in most cases, are strongly active in one tissue and less active in all other tissues (Figure 6). The remarkable exception in this regard were developing (*SMXL5_pro_*) and differentiated (*APL_pro_*) phloem cells. In line with their strong ontogenetic relationship, genes very active in one of the two tissues were often very active in the second one (Figure 6). In contrast, clusters containing genes that were very active in fiber cells were mostly distinct from clusters containing genes whose activity was high in developing vessel elements. Similarly, genes characterizing developing phloem cells (*SMXL5_pro_*) were, by large, distinct from genes characterizing developing xylem cells (*PXY_pro_*). This suggested that, although these cell types partly originate from the same procambial precursors (Shi et al., 2019), they quickly establish very distinct profiles.

**Figure 6.**
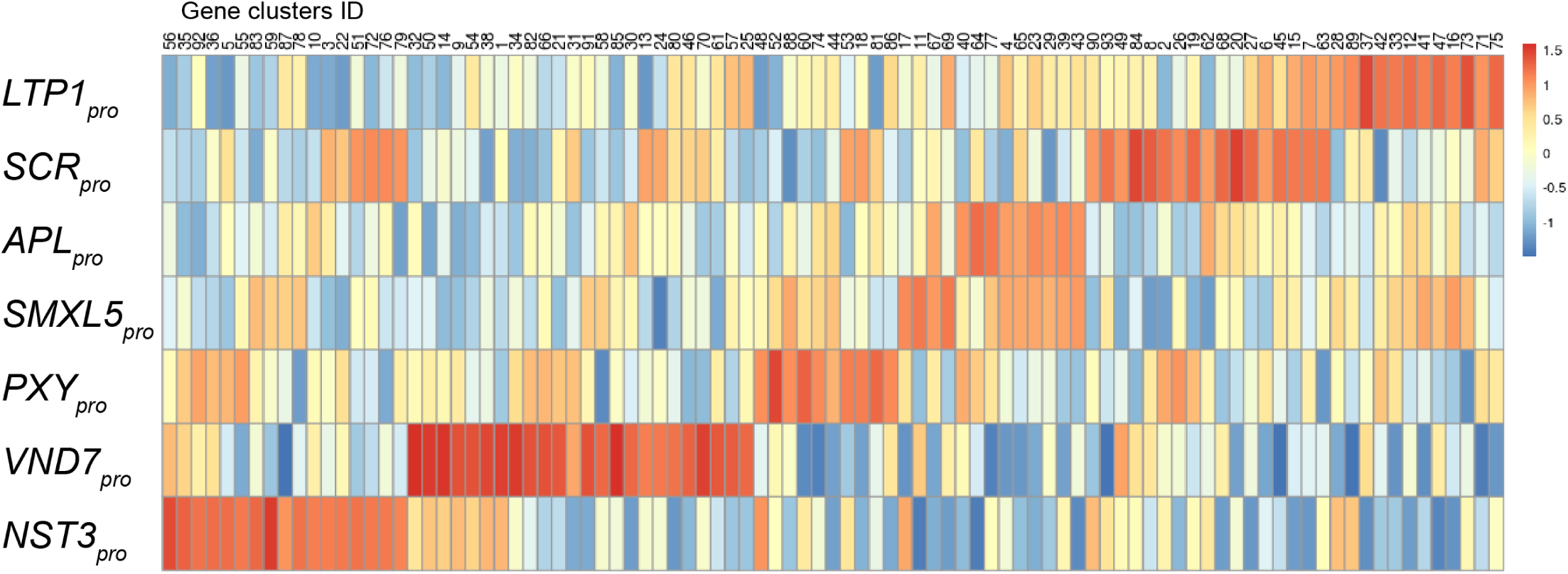
Clustering of significantly differentially expressed genes. Heat map presenting relative gene activities among seven different tissues in each gene cluster. Relative gene expression values are colored according to the color scale on the right. Genes were clustered based on their expression pattern among seven tissues. See Supplemental Figure 5 for more detailed activity profiles of each cluster.

To explore the power of our SDE predictions, we next selected nine genes from the group of genes specifically active in a distinct tissue according to our FANS-derived profiles (Supplemental Dataset 7), generated promoter reporter lines and analyzed their activity in the inflorescence stem (Figure 7G). From the nine promoter reporters generated, three (*AT1G24575_pro_*, *AT1G29520_pro_*, *AT5G28630_pro_*), behaved exactly as predicted (Figures 7C, D, E, F, G, Supplemental Figure 6G, H). *AT1G24575_pro_* and *AT1G29520_pro_* reporters showed activity specifically in the phloem (Figures 7C, D, G; Supplemental Figures 6G, H) and the *AT5G28630_pro_* reporter showed activity specifically in the epidermis (Figures 7E, F). Three other promoter reporters (*AT5G20250_pro_*, *AT5G15970_pro_*, *AT5G16010_pro_*) were active in the predicted tissues but showed also activity in additional tissues, of which most of them, however, were not included in the FANS-based profiling (Figures 7A, B, G, Supplemental Figures 6A, B, I J). The activity of the *AT1G12200_pro_* reporter was, as predicted, observed in phloem cells but not in the cambium possibly due to sensitivity issues (Figure 7G; Supplemental Figure 6E, F). Only two promoter reporters (*AT2G26150_pro_*, *AT5G15870_pro_*) behaved against our predictions with no activity found for the *AT2G26150_pro_* reporter and being active in the phloem instead of the epidermis for the *AT5G15870_pro_* reporter (Figure 7G; Supplemental Figures 6C, D, K, L). Based on these results, and considering that we only included a certain region upstream of the respective start codons into our reporters, we concluded that the predictive power of our FANS/RNA-Seq-based transcriptional profiles was high.

**Figure 7.**
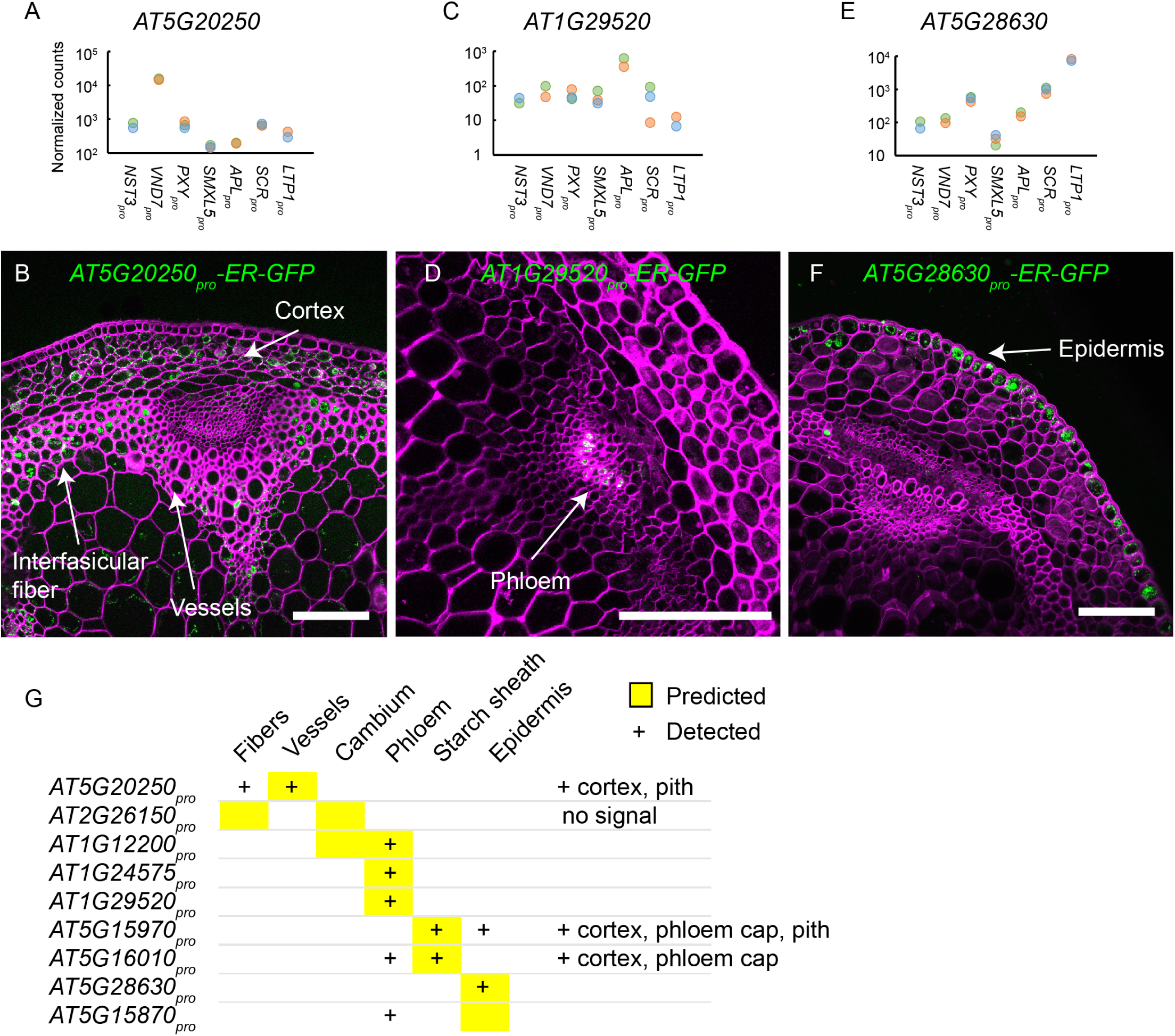
Verification of predicted gene expression patterns. (A, C, E) Normalized read counts among seven different types of nuclei for *AT5G20250, AT1G29520, AT5G28630*. The Y-axis is depicted in logarithmic scale. (B, D, F) Confocal analysis of cross sections from the second internode of promoter reporter lines. The activity of the endoplasmic reticulum (ER)-targeted GFP is shown in green. The cell wall is stained by Direct Red 23 and visualized in magenta. Scale bar: 100 µm. A single focal plane is shown in B and D and a maximum intensity projection is shown in F. At least 2 independent transgenic plant lines for each reporter were observed. (G) Summary of promoter reporter analyses. Yellow background indicates the predicted expression pattern whereas „+“ indicates the observed expression pattern.

SDEs mostly active in the phloem cap were identified by comparing the phloem cap-derived dataset with the dataset derived from the remaining vascular bundle area. This comparison resulted in 575 phloem cap-associated genes (Supplemental Dataset 8, BH adjustment of p value in Wald test < 0.01, fold change >2). As expected (Xu et al., 2019), among this group of genes the GO term ‘glucosinolate biosynthetic process’ (GO:0019758) was enriched (Supplemental Dataset 9). For the pith, we identified 1,633 genes whose reads were significantly enriched in pith-derived samples in comparison to samples from the vascular bundle area (Supplemental Dataset 9, BH adjustment of *p* value in Wald test < 0.01, fold change >2). Performing GO term analyses for this group of genes, identified the term ‘cell death’ (GO:0008219) to be enriched (Supplemental Dataset 9). This was in accordance with previous findings that programed cell death is a characteristic for pith cells (Fujimoto et al., 2018). Taken together, these analyses demonstrated that our LCM-based transcriptomes recapitulated expression profiles of characterized genes and provide informative insights into phloem cap and pith-specific cellular processes.

### Pairwise comparisons of tissues identify tissue-associated gene activities

To see whether other pairwise comparisons were useful for predicting tissue-specific processes, we directly contrasted expression profiles from functionally and ontogenetically related tissues. Fibers (*NST3_pro_*) and xylem vessels (*VND7_pro_*) both determine wood properties in angiosperms and show distinct morphologies which are important for conducting their specific functions (Evert, 2006). Accordingly, their functional specialties are reflected in very distinct expression profiles (Figure 6). Comparing *NST3_pro_*- and *VND7_pro_*-derived datasets, we identified 991 genes as being predominantly expressed in *NST3_pro_*-positive nuclei and 1,503 genes as being predominantly expressed in *VND7_pro_*-positive nuclei (Figure 8A, Supplemental Dataset 10, BH adjustment of *p* value in Wald test < 0.01, fold change >2). Using Gene Ontology (GO) term enrichment analysis (Mi et al., 2019), we found that the terms photosynthesis (GO:0015979), sulfate assimilation (GO:0000103) and response to cytokinin (GO:0009735) and other stimulus-related genes are significantly enriched within the group of *NST3_pro_*-derived genes, whereas the terms xylem vessel differentiation (GO:0048759) and cell wall biogenesis (GO:0042546) were, as expected, enriched within classifications of *VND7_pro_*-derived genes (Figure 8A, Supplemental Dataset 11).

**Figure 8.**
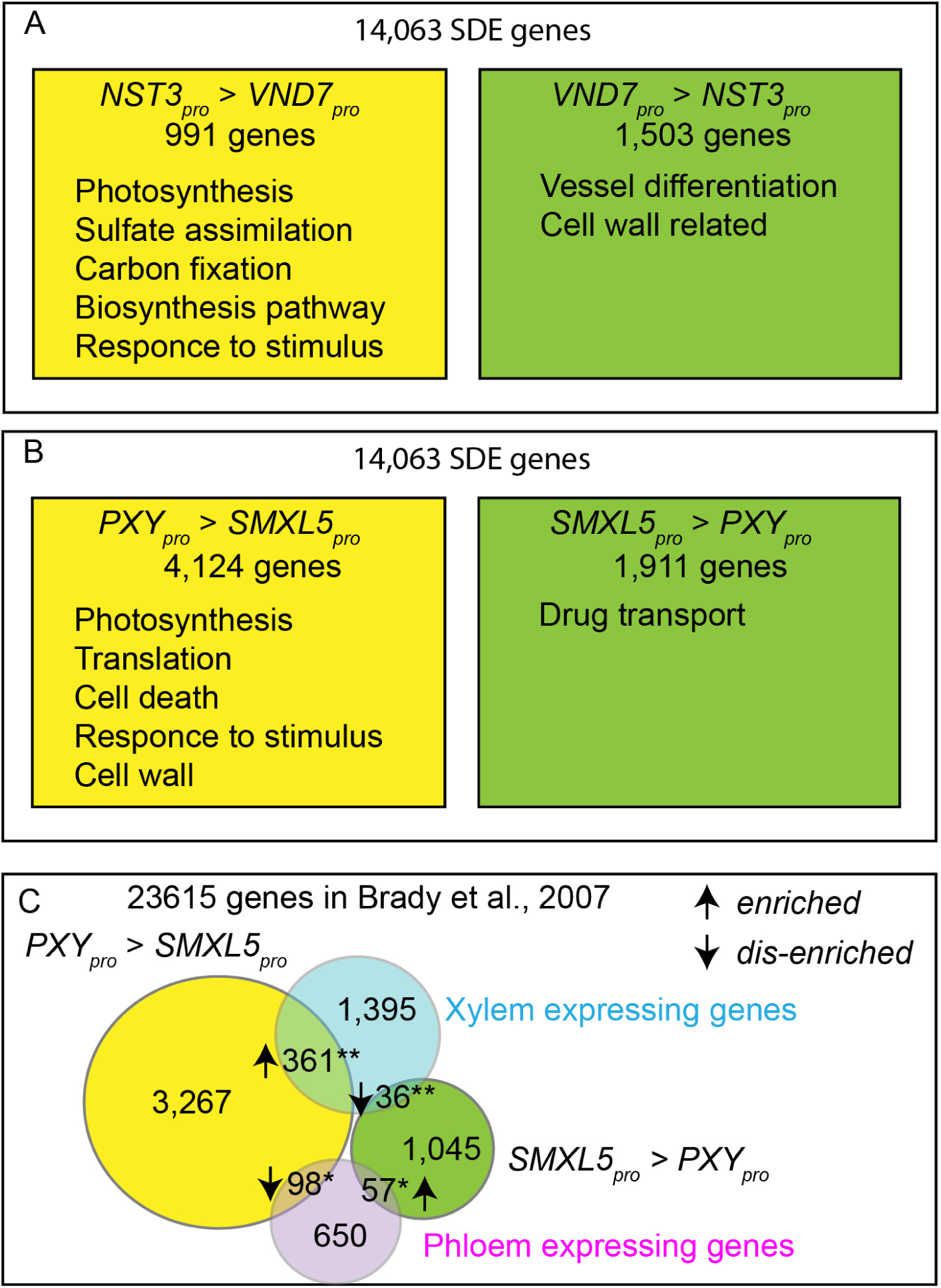
Pairwise comparisons of FANS-derived datasets and comparison with root transcriptome datasets. (A) Summary of pairwise comparison of *NST3_pro_* and *VND7_pro_* datasets (A) and *PXY_pro_* and *SMXL5_pro_* datasets (B). The numbers of SDE genes for each pairwise comparison are shown (BH adjustment of *p* value in Wald test < 0.01, fold change > 2). For detailed GO terms enrichment analysis results, please consult Supplemental Dataset 11 and 13, respectively. (C) Comparison of *PXY_pro_* and *SMXL5_pro_*-associated genes with genes associated in the xylem or phloem of roots. * and ** indicate *p* < 0.01 and *p* < 1e-06 in Fisher’s Exact test, respectively.

Because secondary xylem and phloem tissues originate from cells located in *PXY_pro_* and *SMXL5_pro_* activity domains, respectively (Shi et al., 2019), we also contrasted expression data from those tissues. When directly comparing data from *PXY_pro_* and *SMXL5_pro_*-positive nuclei, we identified 4,124 *PXY_pro_*-associated and 1,911 *SMXL5_pro_*-associated genes (Figure 8B, Supplemental Dataset 12, BH adjustment of *p* value in Wald test < 0.01, fold change >2). Performing GO term enrichment analyses, we found the terms photosynthesis (GO:0015979), response to stimulus (GO:0051869) and translation (GO:0006412) enriched among *PXY_pro_*-associated genes, whereas the term drug transport (GO:0015893) was enriched within the group of *SMXL5_pro_*-associated genes (Figure 8B, Supplemental Dataset 13). Interestingly, genes expressed in the xylem of roots (combined S4, S18, JO121 datasets in (Brady et al., 2007)) were over-represented in the group of *PXY_pro-_*associated genes but under-represented among *SMXL5_pro_*-associated genes (*p* < 1e-06 in Fisher’s Exact Test; Figure 8C). In comparison, genes expressed in the root phloem (combined SUC2, S32, APL, S17 datasets in (Lee et al., 2006; Brady et al., 2007)) were over-represented among *SMXL5_pro_*-associated genes but under-represented among *PXY_pro_*-associated genes (*p* < 0.01 in Fisher’s Exact Test; Figure 8C). This observation suggested that a substantial amount of genes is shared between vascular tissues of primary roots and stems but that there are also large differences in the molecular signatures comparing vascular tissues of both organs.

### Identification of enriched transcription factor binding sites

Networks of transcription factors are vital for establishing tissue-specific gene expression profiles and, thereby, determine cellular behavior (Gaudinier and Brady, 2016). Taking advantage of our identified SDE genes, we therefore sought identifying transcription factor binding regions enriched in promoters of genes with similar expression patterns. To this end, we first assessed significance for the overlap between promoters of genes from each SDE cluster and binding profiles for 387 transcription factors, derived from massive DNA affinity purification sequencing (DAP-Seq) (O’Malley et al., 2016). Among 31 clusters which each contained more than 150 genes, we identified significant enrichment in the overlap with transcription factor binding profiles in 13 clusters (Table 2, Supplemental Dataset 6; *p* < 8.8e-05 (Bonferroni adjusted threshold of 0.05) in Fisher’s exact test). Enrichment for the overlap with transcription factor binding profiles within upstream regions of phloem cap-associated genes identified six putative transcriptional regulators from the NAC, MYB and REM families (Table 2, Supplemental Dataset 14). From them, the poorly studied ANAC028 transcription factor was mostly active in the phloem cap itself, in addition to be expressed in the protophloem of roots (Brady et al., 2007). Transcription factor binding region enrichment analyses for pith-associated genes predicted a set of 61 potential regulators (Table 2, Supplemental Dataset 14). From those, homeodomain transcription factors (ATHBs), PHAVOLUTA (PHV), and the APETALA 2/Ethylene Response Factors (AP2/ERF) members ERF34 and ERF38 were among the genes specifically expressed in the pith.

**Table 2.** Enriched transcription factor (TF) binding regions in SDE clusters derived from FANS and in phloem cap or pith-associated genes derived from LCM. * A part of the whole list is shown. Please refer to Supplemental Dataset 14 for the enrichment fold value and the full list.

| Gene list | Number of genes | Region with high expression | Number of enriched TF binding regions | Associated TFs* |
| --- | --- | --- | --- | --- |
| **FANS cluster ID** |  |  |  |  |
| 8 | 436 | *SCR<sub>pro</sub>* | 52 | CBF1-4; BIM2; CAMTA1, 5; GBF3; ABI5; RTV1; WRKY |
| 7 | 245 | *SCR<sub>pro</sub>, LTP1<sub>pro</sub>* | 41 | CAMTA1, 5; WRKY |
| 19 | 181 | *SCR<sub>pro</sub>, PXY<sub>pro</sub>* | 35 | CAMTA1, 5; WRKY |
| 24 | 358 | *VND7<sub>pro</sub>, SCR<sub>pro</sub>* | 20 | TINY; AREB3; GBF3; bZIPs; ERF38; CAMTA5 |
| 14 | 528 | *VND7<sub>pro</sub>* | 17 | VND1-2; SMB; MYB; GT2 |
| 15 | 223 | *SCR<sub>pro</sub>, LTP1<sub>pro</sub>* | 16 | CBF1-4; WRKY |
| 21 | 220 | *VND7<sub>pro</sub>, PXY<sub>pro</sub>* | 15 | CAMTA1; PHV; LMI1; ATHB |
| 22 | 477 | *NST3<sub>pro</sub>, SCR<sub>pro</sub>* | 4 | ABF2; HY5; MYB55; MYB83 |
| 5 | 445 | *NST3<sub>pro</sub>, PXY<sub>pro</sub>* | 3 | TBP3 |
| 3 | 192 | *NST3<sub>pro</sub>, SCR<sub>pro</sub>* | 1 | ABI5 |
| 38 | 265 | *NST3<sub>pro</sub>, VND7<sub>pro</sub>* | 1 | AT1G49560 |
| 41 | 790 | *LTP1<sub>pro</sub>, SMXL5<sub>pro</sub>* | 1 | ATHB13 |
| 49 | 183 | *SCR<sub>pro</sub>, VND7<sub>pro</sub>* | 1 | ERF38 |
| **LCM** |  |  |  |  |
| Phloem cap | 576 | Phloem cap | 6 | NAC, MYB |
| Pith | 1634 | Pith | 61 | ATHBs; PHV; ERF34,38; Indeterminate-domain (IDD) proteins |

The set of identified factors or their larger families contains promising candidates for determining tissue-specific signatures. In cluster 14, for example, which was associated with developing vessel elements, we found potential binding regions for 17 different transcription factors to be enriched in respective promoters. These factors included VASCULAR-RELATED NAC-DOMAIN1 (VND1, 4.3 fold enrichment), and VND2 (4.6 fold enrichment) (Table 2, Supplemental Dataset 14). Because these transcription factors are expressed in developing vessels where they promote secondary cell wall formation (Zhou et al., 2014), our analysis indeed holds the potential to identify tissue-specific regulators in the inflorescence stem. In turn, the large number of clusters for which no enrichment was detected suggests that, in general, established tissues do not depend on a small set of transcriptional regulators maintaining identity of mature tissues.

## Discussion

Organs as functional units are composed of different tissues determining distinct aspects of their performance. By combining FANS and LCM-based tissue harvesting with RNA-seq analyses, we established a tissue-specific gene expression atlas of the primary Arabidopsis inflorescence stem. In addition to be accessible via the GEO depository (Barrett et al., 2013) under the accession number GSE142034, data for genes of interest can be accessed via a web-site allowing extraction of expression profiles (http://arabidopsis-stem.cos.uni-heidelberg.de/).

Several observations underline the robustness of our mRNA profiling. First, the activities of six out of seven genes whose promoters were used for tissue-specific nucleus labeling were found to peak in the respective tissues. This observation suggests that the chosen promoters mostly recapitulate the expression patterns of endogenous genes. Moreover, nuclei keep a sufficient amount of their mRNA content during the isolation process and the contamination by cytosolic mRNA, which is released during tissue disruption, is apparently low. The fact that *LTP1* activity did not peak in *LTP1_pro_*-positive nuclei, may reveal discrepancies between the activity pattern of our chosen *LTP1* promoter and the distribution of the endogenous *LTP1* mRNA. It is also important to be aware that the stability or movement of the *LTP1* mRNA and the *H4-GFP* mRNA driven by the *LTP1* promoter may be very different. This may result in discrepancies between patterns of H4-GFP protein accumulation and *LTP1* mRNA accumulation. However, our comparative analyses suggests that we succeeded in isolating epidermis-specific mRNA.

The second observation supporting robustness of obtained profiles is that promoters from genes we predicted to have a distinct spatial pattern recapitulated, by large, these patterns. In addition to the possible reasons resulting in differences between the accumulation of endogenous transcripts and the H4-GFP protein discussed above, it is important to note that the promoter regions upstream of respective start codons, which we chose to drive fluorescent reporters, may miss regulatory elements substantially influencing gene expression of endogenous genes (Raatz et al., 2011). Therefore, a certain level of differences between FANS/LCM-derived profiles and reporter activities are expected and do not necessarily argue for a low predictive power of our transcriptional profiles. As a third observation arguing for the relevance of our profiles, the activity pattern of genes and pathways known to be associated with certain tissues, was reflected in our datasets. Taken together, we conclude that the obtained expression data provide a realistic picture of gene activities in the Arabidopsis inflorescence stem.

Although cytosolic and nuclear mRNA populations may differ due to a tight regulation of nuclear export or differences in mRNA homeostasis (Yang et al., 2017; Choudury et al., 2019), our results indicate that gene expression profiles of mature plant organs can be faithfully characterized by profiling nuclear mRNA. Interestingly, our RNA-seq results show that the sensitivity of FANS/RNA-seq-based analyses is comparable or even higher than conventional profiling of whole organs by RNA-seq (Table 1). In comparison to the profiling of mRNA from whole cells, additional advantages of fluorescence-based profiling of nuclear mRNA are important to consider (Lake et al., 2017; Abdelmoez et al., 2018; Bakken et al., 2018; Wu et al., 2019). First, profiling nuclear mRNA allows analyzing differentiated plant tissues with prominent cell walls or heterogenous sizes of protoplasts which can be labeled by genetically encoded fluorescent reporters. Thus, the method allows targeting a broad range of tissues independent from enzymatic accessibility or morphological markers which are required for tissue identification when using LCM. Second, following our procedure, tissue disruption and subsequent washing steps during nucleus extraction takes 30 to 40 minutes. Depending on cell wall properties, protoplasting often requires more time (Birnbaum et al., 2003) which increases the risk of a treatment-induced change of transcript abundance. Third, nuclear mRNAs carry a higher ratio of unprocessed mRNA molecules compared to mRNA from whole cells (Lake et al., 2017; Abdelmoez et al., 2018; Wu et al., 2019). Due to a predictable rate of mRNA processing in nuclei, this feature can be used to calculate actual transcription rates of individual genes (Gaidatzis et al., 2015; Lake et al., 2016). Therefore, nuclear RNA-seq datasets provide a different quality of information compared to RNA-seq datasets from whole cells. Fourth, in comparison to precipitation-based methods of nucleus isolation like INTACT (Deal and Henikoff, 2010), thresholds for fluorescence-based nucleus sorting can be adequately set according to the fluorescence intensity. This feature not only provides flexibility in selecting distinct nuclei populations according to the level of reporter activity. We envisage that it also allows combining a multitude of different fluorescent markers to collect distinct and highly specific nucleus populations from single plant lines.

Considering the importance of plant stems and their tissues in land plant evolution on the one side (Xu et al., 2014) and biomass accumulation on the other side (Yamaguchi and Demura, 2010), information on their specific gene expression profiles is certainly vital. Our datasets allow the formulation of testable hypotheses about the activity of distinct pathways in individual stem tissues and their role in determining plant performance overall.

## Methods

### Plant material

*Arabidopsis thaliana* (L.) Heynh. Col-0 plants were used as a genetic background. Plants were grown in short-day conditions (10 h light and14 h darkness) for 3-5 weeks and then transferred to long-day conditions (16 h light and 8 h darkness) for 3 weeks to induce reproductive growth. The *SMXL5_pro_:H4-GFP* (*pIL53*) line and the *PXY_pro_:H4-GFP* (*pPS24*) lines were described previously (Shi et al., 2019). Other transgenic lines were generated through the floral dipping method using *Agrobacterium tumefaciens* (Clough and Bent, 1998). Analyses were performed using homozygous lines except for promoter reporter line analyses (Figure 7 and Supplemental Figure 6) where T1 generation plants were used.

### DNA vector construction

An *H4-GFP*-containing construct was a gift from Daniel Schubert (Freie Universität Berlin, Germany). *NST3_pro_:H4-GFP* (*pMS59*), *VND7_pro_:H4-GFP* (*pTOM61*), *APL_pro_:H4-GFP* (*pPS02*), *SCR_pro_:H4-GFP* (*pPS20*) and *LTP1_pro_:H4-GFP* (*pPS16*) were cloned using the pGreen0229 vector (Hellens et al., 2000) as a backbone. Cloning of *APL* promoter and terminator regions was described previously (Sehr et al., 2010). Primer sequences to clone promoter and terminator regions of other genes and to amplify H4-GFP sequence with appropriate sequence for restriction enzyme are listed in Supplemental Table 1. Constructs for promoter reporters were generated using the GreenGate system (Lampropoulos et al., 2013). Promoter regions were amplified using primers listed in Supplemental Table 1, cloned into the *pGGA000* vector and used for GreenGate reactions with the ER signal peptide sequence (*pGGB006*), the *GFP* sequence (*pGGC014*), the *HDEL* sequence (*pGGD008*), the *RBCS* terminator sequence (*pGGE001*), the BASTA resistance sequence (*pGGF001*) and the destination vector (*pGGZ003*) (Lampropoulos et al., 2013).

### Confocal Microscopy

Free-hand cross sections of the second bottom-most internode of Arabidopsis inflorescence stems were generated using razor blades (Wilkinson Sword, High Wycombe, U.K.) and stained with 0.1 % solution of Direct Red 23 (Hoch et al., 2005; Anderson et al., 2010; Ursache et al., 2018) (30 % content powder, Sigma-Aldrich, St. Louis, U.S.A., #212490) in Ca^2+^, Mg^2+^-free PBS. Sections were then briefly washed using tap water and put into a glass-bottom dish (ibidi, Gräfelfing, Germany, µ-Dish 35 mm, high, 81151). Images were captured using a Nikon A1 confocal microscope using a 25x water immersion objective lens (Nikon, Tokyo, Japan, Apo 25xW MP, 77220) and gallium arsenide phosphide (GaAsP) detectors. GFP and Direct Red 23 were excited by 488 nm and 561 nm lasers, respectively.

### Nucleus isolation

Second internodes of Arabidopsis inflorescence stem were collected in a petri dish on ice. 2 ml cold isolation buffer (20 mM Tris (pH=7.5), 40 mM NaCl, 90 mM KCl, 2 mM Ethylenediaminetetraacetic acid (EDTA) (pH=8.0), 0.5 mM EGTA (ethylene glycol-bis(β-aminoethyl ether)-N,N,N’,N’-tetraacetic acid) (EGTA), 0.05 % Triton X-100, 0.5 mM Spermidine, 0.2 mM Spermine, 15 mM 2-Mercaptoethanol, 0.5 mM Phenylmethylsulfonylfluorid (PMSF), cOmplete Protease Inhibitor Cocktail (Roche, Basel, Switzerland, #11697498001)) supplemented with 10 µl RiboLock RNase inhibitor (40 U/µL) (ThermoFisher, Waltham, U.S.A., #EO0381) was added to 2 g of stem tissue. Samples were chopped manually on ice for 10 min, then transferred to a low binding tube (Protein LoBind Tube, Eppendorf, Hamburg, Germany, #0030 108.132) through a filter (CellTrics 50 µm, Sysmex, Kobe, Japan, # 04-004-2327). Nuclei were collected by centrifugation (1000g, 10min, 4°C) and gently washed two times with cold resuspension buffer (isolation buffer without Triton X-100). 300 µl of resuspension buffer supplemented with 10 µg/ml Hoechst 33342 at final concentration and 5 µl RiboLock RNase inhibitor were added to the collected nuclei, then transferred to a tube through a filter (FALCON Round-Bottom Tube with Cell-Strainer Cap, Corning, New York, U.S.A., #352235) for sorting.

### Nucleus sorting

Nucleus sorting was performed on a BD FACSAria^TM^ IIIu cell sorter (Becton Dickinson, Franklin Lakes, U.S.A.) using a 70 µm sort nozzle. A sheath pressure of 70 psi and a drop drive frequency of 87 kHz was applied. GFP fluorescence was excited at 488 nm using a 20 mW blue laser and Hoechst fluorescence at 405 nm using a 50 mW violet laser. A 530/30 bandpass filter was used for GFP and a 450/40 bandpass filter for Hoechst detection. Autofluorescence at 405 nm excitation was detected using a 585/42 bandpass filter. FSC (forward scatter) detector voltage was set to 55 and a 1.0 neutral density filter was used. SSC (side scatter) detector voltage was set to 275, Hoechst detector voltage to 352 and GFP detector voltage to 379. Events were triggered on FSC using a threshold of 5,000 and Hoechst with a threshold of 400. All the events were first filtered by FSC and SSC (Supplemental Fig 2E, gate 1). Nuclei were then identified based on Hoechst fluorescence. Doublets and aggregates were characterized by their higher Hoechst signal width values and were excluded from sorting by plotting Hoechst signal width against Hoechst signal area (Supplemental Figure 2F, gate 2). To exclude false positive events due to autofluorescence in the yellow-green spectral range, the GFP detection channel was plotted against a yellow fluorescence detection channel and only events with low yellow signal intensity were selected for sorting (Supplemental Figure 2G, gate 3). Then, the gate for GFP positive nuclei selection were set using wild type plants as a reference (Supplemental Figure 2H, gate 4). DNA content distribution was used to monitor the whole procedure (Supplemental Figure 2I).

### RNA extraction, cDNA library preparation and next-generation sequencing

15,000 nuclei for each population were sorted into 750 µl TRIzol reagent (ThermoFisher, Waltham, U.S.A., #15596018), except for *LTP1_pro_*-S21 where 10,800 nuclei were sorted. RNA was precipitated and washed according to the manufacture’s protocol, and then resuspended in 15 µl water. 1 µl of the total RNA was used for mRNA-seq library construction using the Smart-seq2 protocol (Picelli et al., 2013; Hofmann et al., 2019). The resulting cDNA was quantified using the Qubit Fluorometer (Thermo Fisher, Waltham, U.S.A.) and the dsDNA High Sensitivity Assay (Thermo Fisher, Waltham, U.S.A.). DNA quality was checked on the Agilent Bioanalyzer (Agilent, Santa Clara, U.S.A.). Libraries for next generation sequencing were generated using the NEBNext Ultra II gDNA prep kit with the NEBNext Multiplex Oligos for Illumina (New England Biolabs, Ipswich, U.S.A.). Prior to library generation, the samples were fragmented using the Covaris S2 system (Covaris, Woburn, U.S.A.). Libraries were sequenced in the single-end 50 base mode on an Illumina HiSeq 2500 machine (Illumina, San Diego, U.S.A.). When more than three samples were prepared, amplified cDNA samples were subjected to PCR analysis amplifying the *H4-GFP* sequence using (*H4GFPfor4* and *GFPrev3* primers; see Supplemental Table 1 for sequence). The preparations showing high contrast between GFP-positive and GFP-negative samples were chosen for subsequent RNA-seq analysis.

### Laser capture microdissection

LCM, subsequent RNA extraction and amplification was carried out as previously described (Agusti et al., 2011). Library preparation and RNA sequencing were carried out by BGI Genomics (Shenzhen, China) using HiSeq 2000 (Illumina, San Diego, U.S.A.) using the single-end mode 50 base mode.

### Bioinformatic analyses

The TAIR10 genome sequence was obtained through Ensemble Plants (Arabidopsis_thaliana.TAIR10.28.dna.toplevel.fa) (Arabidopsis Genome, 2000; Bolser et al., 2016). The Araport11 gene annotation file was taken (Araport11_GFF3_genes_transposons.201606.gtf) (Cheng et al., 2017), and the chromosome names were changed accordingly to fit the TAIR10 genome dataset. The genome index was generated using STAR (v2.5.0a) (Dobin et al., 2013), and FASTQ files were trimmed by Cutadapt (v2.3) (Martin, 2011) using the following options: *-g AAGCAGTGGTATCAACGCAGAGTACGGG-a CCCGTACTCTGCGTTGATACCACTGCTT -g AAGCAGTGGTATCAACGCAGAGTAC - a GTACTCTGCGTTGATACCACTGCTT -b “A 30” -b “T 30” -n 2 -g AGATCGGAAGAGC -l 50 -m 23 --overlap 5.* By these settings, oligo sequences used for the Smart-seq2 amplification, poly A or T sequences, and Illumina adaptor sequences were removed from the reads. Trimmed files were then mapped to the genome by STAR using *--outFilterMultimapNmax 1 --alignIntronMax 10000 -- alignMatesGapMax 10000 --outFilterScoreMinOverLread 0.9* options. Intron length limit (10,000 bp) were set based on the characteristics of the Arabidopsis genome (Chang et al., 2017). Please consult Supplemental Table 2 for basic statistics applied on RNA-seq datasets. For LCM-derived RNA-seq datasets, untrimmed reads were mapped with same settings. For determining GFP reads, the GFP sequence was indexed by STAR using *–genomeSAindexNbases* 4 option. Subsequently, reads were mapped using STAR using the *--alignIntronMax 1 --alignMatesGapMax 1 -- outFilterScoreMinOverLread 0.90* options. Further analysis was carried out in R (v3.5.0). The GTF file was imported using makeTxDbFromGRanges in GenomicFeatures library (v1.32.3) with *drop.stop.codons* option (Lawrence et al., 2013). The position of each gene was extracted by *genes* function, which also includes the intron region. Read counts per gene were obtained using summarizeOverlaps in GenomicAlignments library (v1.16.0) using the *mode = “Union”, ignore.strand = TRUE* options (Lawrence et al., 2013). For TPM calculation, the *width* function was used to calculate the length of each gene extracted, which also included intron regions. For the analysis of intron and exon regions, *intronsByTranscript* and *transcripts* functions were used for genomic range extraction. For analyzing intergenic regions, the *gaps* function was used with the extracted gene regions (output of *genes* function) for genomic range extraction. For comparison with previous datasets, three replicates of the Mock_bdl dataset (deposited in NCBI’s Gene Expression Omnibus (GEO) database (Barrett et al., 2013): GSE98193) were used (Brackmann et al., 2018). Differentially expressed genes were identified using default options of DESeq2 (v1.20.0) (Wald test) for comparing GFP-positive and GFP-negative nucleus populations, and the LRT in DESeq2 was used for multiple comparison (Love et al., 2014). For GFP reads, the inverted beta-binomial test was performed by the *ibb* function (Pham and Jimenez, 2012), using the sum of uniquely mapped reads and multiple mapped reads to the Arabidopsis genome as the total sample count. PCA analyses were carried out by the *plotPCA* function in R using log transformed values obtained from the *rlog* function in the DESeq2 library with *blind=FALSE* options. Distances between samples were calculated by *dist* function in *stats* library (3.5.0) and visualized in heat map by *pheatmap*. GO enrichment analysis was carried out using PANTHER (v14.1) through TAIR10 platform (https://www.arabidopsis.org/tools/go_term_enrichment.jsp) (Mi et al., 2019). Fisher’s Exact test was carried out in R. Basic statistics were carried out by Microsoft Excel.

### Determining enrichment of transcription factor binding regions

For determining the enrichment of transcription factor binding regions, 568 genome-wide DAP-Seq profiles for 387 Arabidopsis TFs were downloaded from the Plant Cistrome Database (O’Malley et al., 2016). The foreground dataset was taken as respective (−1500;+1) 5’-regions of the SDE genes revealed by FANS/RNA-seq or LCM/RNA-seq experiments; the background dataset compiled as 5’-regions upstream to start codons of 19916 protein-coding genes from TAIR10 version of the Arabidopsis genome (Lamesch et al., 2012). To assess the significance of overlaps between the 5’-regions and the transcription fac tor binding profiles, we counted the numbers of foreground/background 5’-regions that contained at least 1 bp overlap with each of the DAP-Seq profiles and applied Fisher’s Exact Test. A transcription factor was considered as a potential regulator of the gene set if its binding profile was significantly more often represented in the 5’-regions of the respective genes compared to the rest of the genome under Bonferroni adjusted *p* < 8.8e-05.

### Accessing RNA sequencing data

Raw data discussed in this publication have been deposited in NCBI’s Gene Expression Omnibus (Barrett et al., 2013) and are accessible through GEO Series accession number GSE142034 at https://www.ncbi.nlm.nih.gov/geo/. In addition, data for genes of interest can be accessed via a web-site based tool allowing extraction of expression profiles (http://arabidopsis-stem.cos.uni-heidelberg.de/).

### Accession Numbers

Arabidopsis Genome Initiative locus identifiers for each genes are: *NST3 (AT1G32770), VND7 (AT1G71930), PXY (AT5G61480), SMXL5 (AT5G57130), APL (AT1G79430), SCR (AT3G54220), LTP1 (AT2G38540), H4 (AT5G59690), XCP1 (AT4G35350), WOX4 (AT1G46480), MOL1 (AT5G51350), PEAR1 (AT2G37590), PEAR2 (AT5G02460), NAC086 (AT5G17260), PIN3 (AT1G70940), FDH (AT2G26250), CER6 (AT1G68530), NST1 (AT2G46770), VND6 (AT5G62380), SEOR1 (AT3G01680), VND1 (AT2G18060), VND2 (AT4G36160; NAC076), ABI5 (AT2G36270), CAMTA1 (AT5G09410), CAMTA5 (AT4G16150), CBF1 (AT4G25490), CBF2 (AT4G25470), CBF3 (AT4G25480; DREB1A), CBF4 (AT5G51990), GBF3 (AT2G46270), RTV1 (AT1G49480), SMB (AT1G79580), GT2 (AT1G76890), PHV (AT1G30490), LMI1 (AT5G03790), MYB55 (AT4G01680), MYB83 (AT3G08500), ABF2 (AT1G45249), HY5 (AT5G11260), TINY (AT5G25810), AREB3 (AT3G56850), ATHB13 (AT1G69780), ERF38 (AT2G35700), ERF34 (AT2G44940), CYP83A1 (AT4G13770), CYP83B1 (AT4G31500), ANAC028 (AT1G65910), ANAC074 (AT4G28530)*.

## Supplemental Data

**Supplemental Table 1.** Primers used in this study.

**Supplemental Table 2.** Basic statistics of RNA-seq dataset in this study.

**Supplemental Figure 1.** H4-GFP reporter lines used in this study.

**Supplemental Figure 2.** FANS for GFP-positive and GFP-negative nuclei.

**Supplemental Figure 3.** PCA plot and correlation heatmap of all datasets derived from GFP-positive nuclei of seven different reporter lines.

**Supplemental Figure 4.** Gene expression profiles for epidermis-associated genes.

**Supplemental Figure 5.** Gene expression profiles for each cluster of genes.

**Supplemental Figure 6.** Additional promoter reporter lines analysis.

**Supplemental Dataset 1.** Significantly differentially expressed genes in transcriptome datasets obtained from GFP-positive comparing to GFP-negative nuclei from *NST3_pro_*, *SMXL5_pro_* and *APL_pro_* lines, respectively.

**Supplemental Dataset 2.** Raw read counts for each RNA-seq dataset.

**Supplemental Dataset 3.** Raw and normalized read counts for each LCM-derived RNA-seq dataset.

**Supplemental Dataset 4.** The result of LRT analysis of RNA-seq datasets obtained from *NST3_pro_, VND7_pro_, PXY_pro_, SMXL5_pro_, APL_pro_, SCR_pro_* and *LTP1_pro_*-positive nuclei.

**Supplemental Dataset 5.** Normalized read counts for each RNA-seq dataset.

**Supplemental Dataset 6.** Clustering of genes based on their expression pattern among seven tissues.

**Supplemental Dataset 7.** Average value of normalized read counts of SDE genes within each nuclei type, ranked by the values.

**Supplemental Dataset 8.** SDE genes comparing Phloem cap/Pith and the remaining vascular bundle in LCM-derived datasets.

**Supplemental Dataset 9.** GO term enrichment analysis in phloem cap-associated or pith-associated genes.

**Supplemental Dataset 10.** SDE genes comparing *NST3_pro_*-positive and *VND7_pro_*-positive nuclei.

**Supplemental Dataset 11.** GO term enrichment analysis among genes specifically active in *NST3_pro_*-positive when compared to *VND7_pro_*-positive nuclei, and among genes specifically active in *VND7_pro_*-positive nuclei compared to *NST3_pro_*-positive nuclei.

**Supplemental Dataset 12.** SDE genes comparing *PXY_pro_*-positive nuclei and *SMXL5_pro_*-positive nuclei.

**Supplemental Dataset 13.** GO term enrichment analysis among genes specifically active in *PXY_pro_*-positive when compared to *SMXL5_pro_*-positive nuclei, and among genes specifically active in *SMXL5_pro_*-positive when compared to *PXY_pro_*-positive nuclei.

**Supplemental Dataset 14.** Fold enrichment values of significantly enriched transcription factor binding regions in the upstream regions of genes from FANS/RNA-seq-derived clusters and tissue-specific genes from LCM/RNA-seq analyses.

## Acknowledgments

We would like to thank Daniel Schubert (Freie Universität Berlin, Germany) for providing reagents, Michael Nodine (GMI, Vienna, Austria) for providing RNA amplification protocols, Florent Murat (ZMBH, Heidelberg University, Germany) and Dmitry Oschepkov (Institute of Cytology and Genetics SB RAS, Russia) for providing advice on bioinformatics analysis. We also thank Jan U. Lohmann (COS, Heidelberg University, Germany) for providing instruments. Library preparation and next generation sequencing was carried out by David Ibberson (The CellNetworks Deep Sequencing Core Facility, Heidelberg University, Germany). Nucleus sorting was carried out by Monika Langlotz (Flow Cytometry and FACS Core Facility, ZMBH, Heidelberg University, Germany). We also thank Ivan Lebovka, Min-Hao Chiang, Ilona Jung, Nihad Softic (COS, Heidelberg University, Germany) and Martina Laaber-Schwarz (GMI, Vienna, Austria) for technical assistance and Steffen Greiner (COS, Heidelberg University, Germany) for comments on the manuscript. This work was supported by the European Research Council through an ERC Consolidator grant [PLANTSTEMS to T.G.], the Deutsche Forschungsgemeinschaft [DFG, through a Heisenberg Professorship GR2104/5-2 and the SFB873 to T.G.], the Alexander von Humboldt-Stiftung [3.5-JPN −1164674-HFST-P to D.S.], the European Molecular Biology Organization [ALTF 342-2012 to V.J.], the Russian Foundation for Basic Research and the DFG [RFBR-DFG 19-54-12013 for cooperation of Russian and German teams] and the Russian Budget project [0324-2019-0040 to V.M and V.L].

## Author Contributions

T.G. designed the research. D.S., V.J., J.A., V.K., V.M. and P.S. performed the research. P.S. and V.K. generated plant material. V.J. and P.S. developed nucleus isolation and RNA extraction protocols. V.L. and V.M. developed the method for enrichment analysis of transcription factor binding regions. D.S., V.J., V.L. and V.M. analyzed data. D.S. and T.G. wrote the paper.

