## Supplemental Data for "Tissue-specific transcriptome profiling of the *Arabidopsis thaliana* inflorescence stem reveals local cellular signatures"

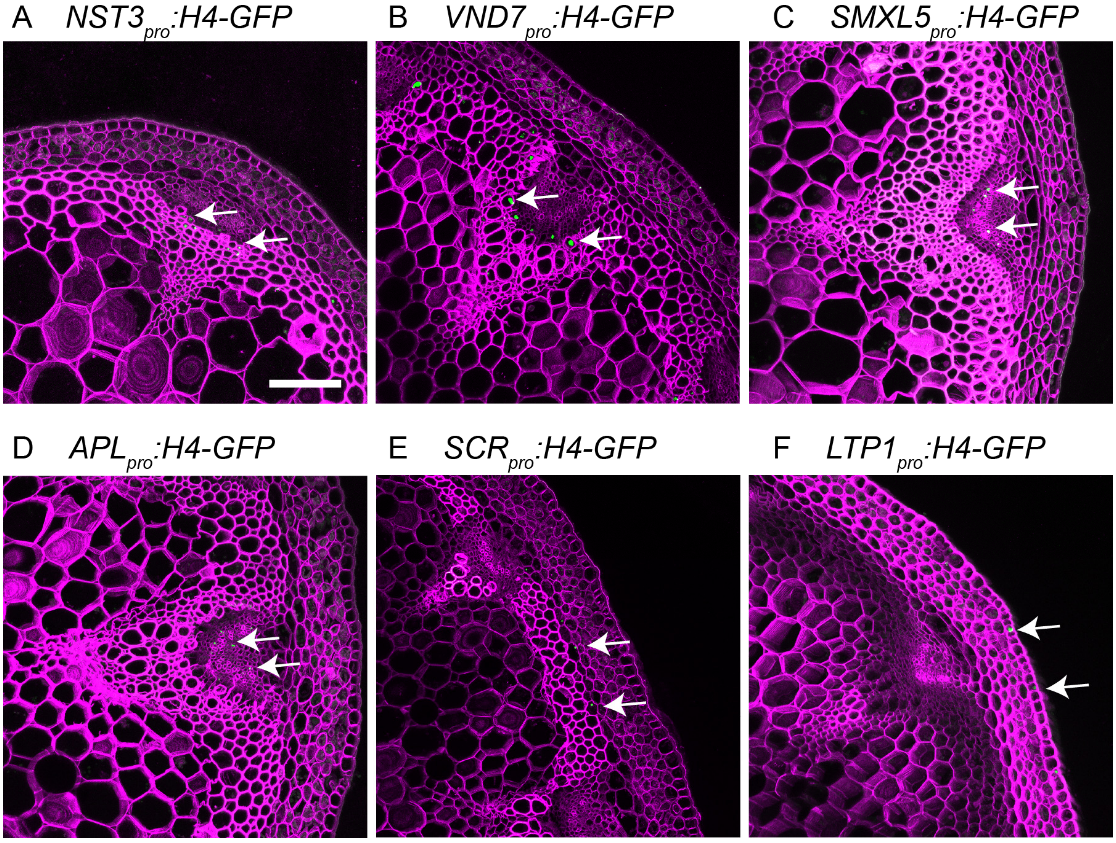

**Supplemental Figure 1. H4-GFP reporter lines used in this study (supports Figure 1)**

(A-F) Maximum intensity projection of confocal images of cross sections from the second stem internode. GFP signal is shown in green and cell walls are stained by Direct Red 23 and visualized in magenta. Arrows indicate GFP-positive nuclei. Scale bar: 100 µm. Note that only the nuclei in the limited observable depth of the section are detected and visualized.

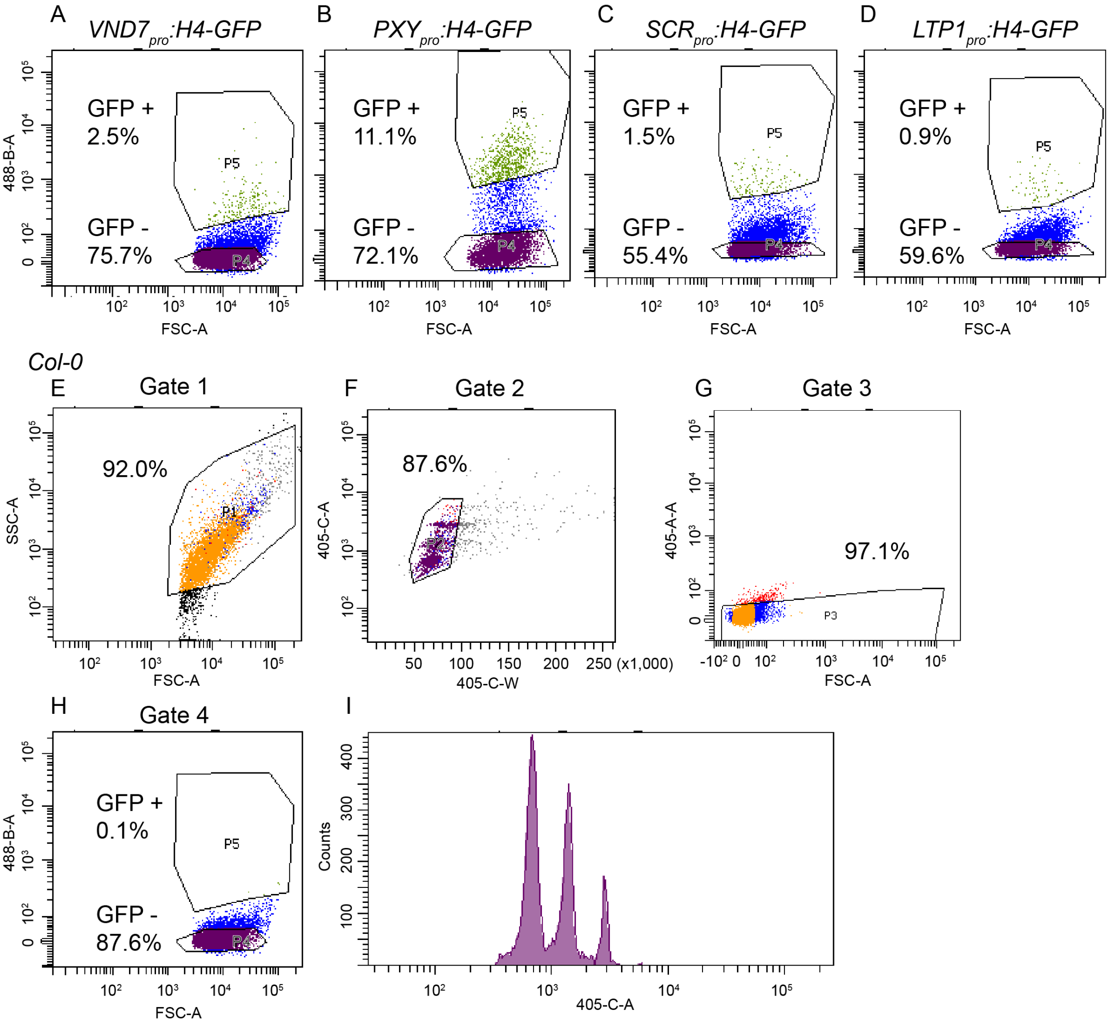

**Supplemental Figure 2. FANS for GFP positive and negative nuclei (supports Figure 2)**

訃悉-D) Plot of gate settings defining GFP-positive (P5) and GFP-negative nucleus (P4) populations from *VND7$_{pro}$:H4-GFP* (A), *PXY$_{pro}$:H4-GFP* (B), *SCR$_{pro}$:H4-GFP* (C) and *LTP1$_{pro}$:H4-GFP* (D) transgenic plants, respectively. The ratio of each population compared to all events are indicated. X axis (FSC; forward scatter intensity) corresponds to the diameter of the cell, the Y axis (488) indicates levels of GFP fluorescence. (E-H) Additional FANS gate settings. One example from wild type plants without transgene sorted in parallel to samples from *VND7$_{pro}$:H4-GFP* plants is shown. See Nucleus Sorting section in the Methods section for further description. SSC: side scatter (E), 405-C-W: width of fluorescent signal induced by the 405 nm laser, 405-C-A: area of fluorescent signal induced by the 405 nm laser (F), 405-A-A : auto-fluorescence signal detected in the YFP channel excited by the 405 nm laser (G). (H) Same gate settings used in (A). (I) Histogram showing the distribution of DNA content of nuclei detected by 405 laser after Hoechst staining.

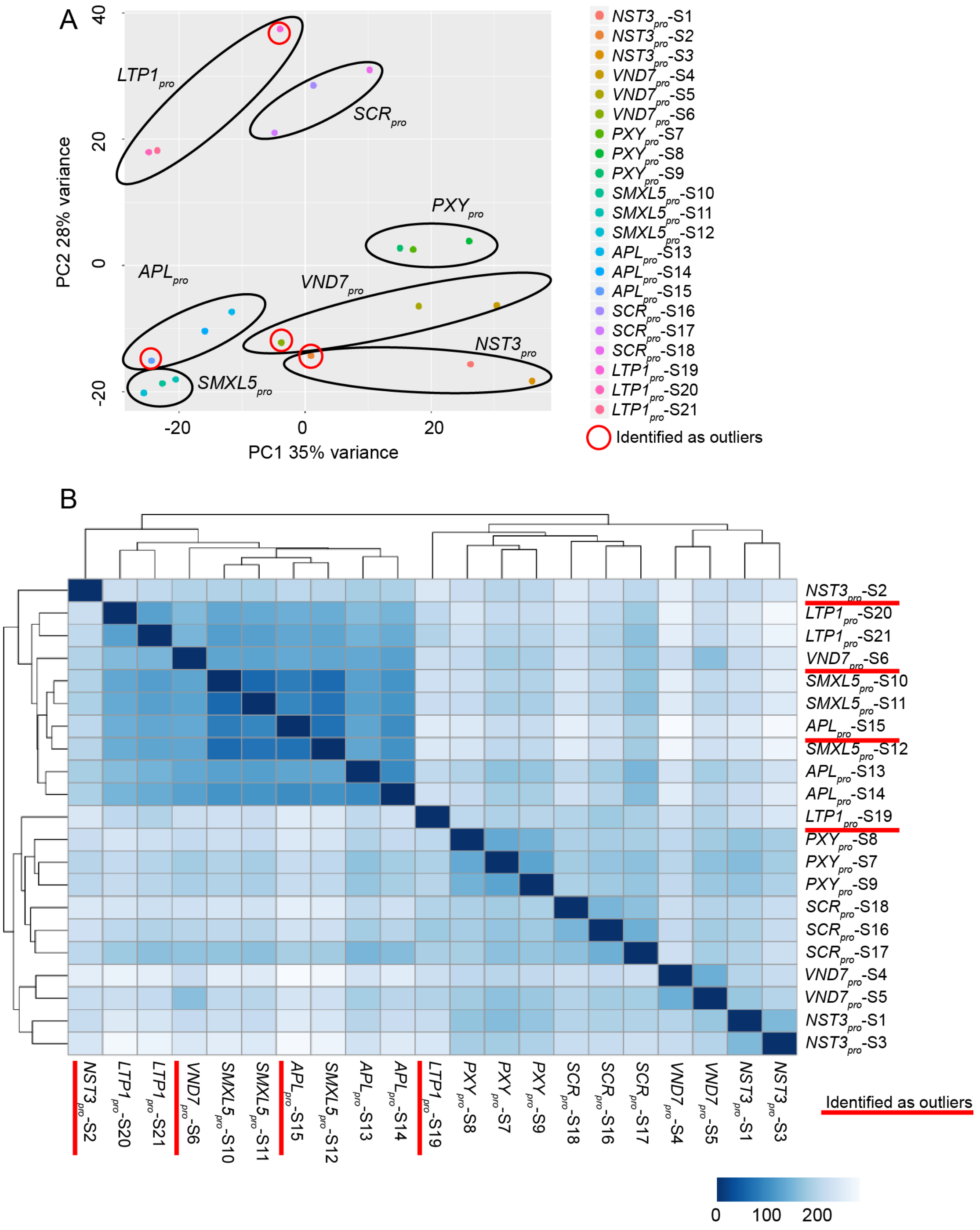

**Supplemental Figure 3. PCA plot and correlation heatmap of all datasets derived from GFP-positive nuclei of seven tissues (supports Figure 3).**

(A) Principal component analysis (PCA) on log-transformed normalized read counts of each RNA-seq dataset. (B) Heatmap indicating statistical distances between RNA-seq dataset according to the color code. Three replicates were obtained from each nucleus population (seven different promoter lines). Samples marked in red did not cluster together with other replicates of the same nucleus population.

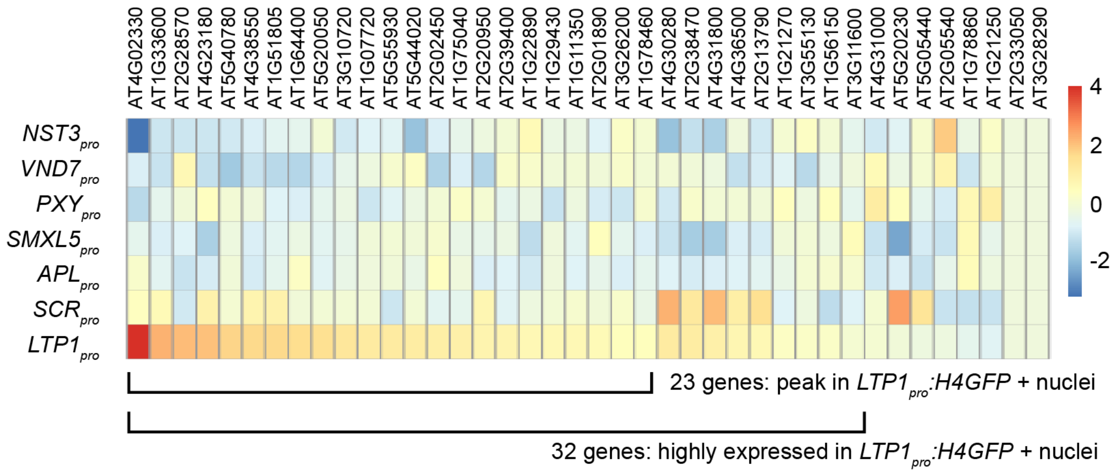

**Supplemental Figure 4. Gene expression profiles for epidermis-associated genes (supports Figure 4).**

Relative expression heat map of the 40 genes most specifically expressed in the epidermis according to Suh et al., 2005 in seven FANS-derived datasets. Relative expression values are color coded indicating the log2 values of normalized read counts after the mean values found in all seven nucleus samples for each gene were subtracted.

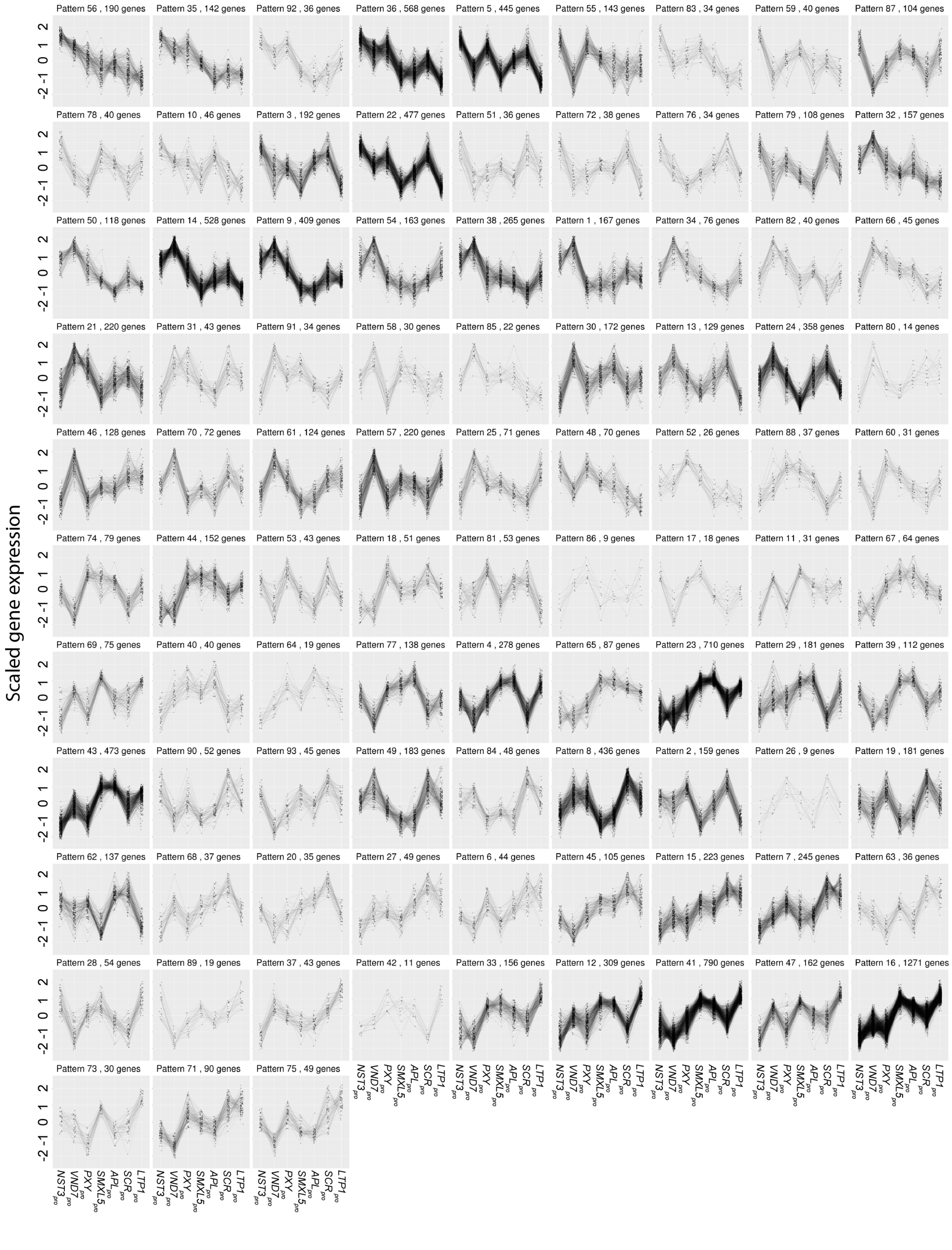

**Supplemental Figure 5. Gene expression profiles for each FANS-derived gene cluster (supports Figure 6).**

Scaled relative gene expression values are visualized for each cluster. Normalized read counts were log2-transformed and then the distribution range of each gene among 7 nucleus type were again normalized. The pattern of expression is ordered in the same way as displayed in Figure 6.

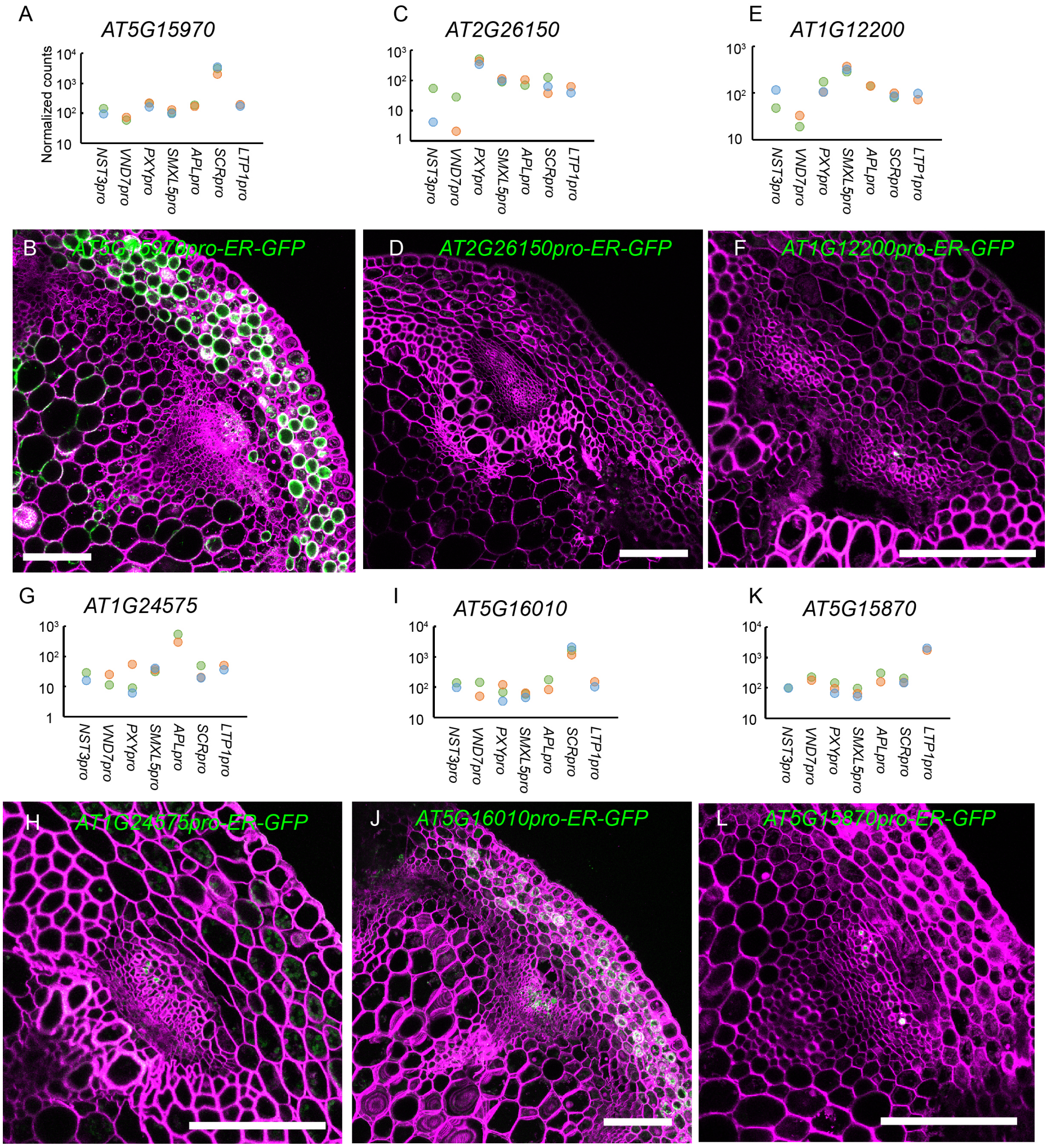

**Supplemental Figure 6 Additional promoter reporter lines analysis. (supports Figure 7)**

(A, C, E, G, I, K) Normalized gene read counts among seven different tissues for *AT5G15970, AT2G26150, AT1G12200, AT1G24575, AT5G16010, AT5G15870*. The Y-axis is in the logarithmic scale. (B, D, F, H, J, L) Confocal images of cross sections from the second internode of different promoter reporter lines. The signal from the endoplasmic reticulum (ER)-targeted GFP protein is shown in green and cell walls are stained by Direct Red 23 and visualized in magenta. Scale bars: 100 µm. A single focal plane is shown in B, D, F, H, L and a maximum intensity projection is shown in J. At least two independent transgenic plant lines for each transgene were analyzed.

**Supplemental Table 1. Primers used in this study.**

| Primer name | Sequence (5‘-3‘) | Usage |
|---|---|---|
| *NST3for3* | ACTAGCGGCCGCgattctacacattcacaaagtttactac | NST3 promoter |
| *NST3rev4* | GTCGACTGAGATCTAGCCATGGttaacgaagatagc aatatattttggg | NST3 promoter |
| *NST3for5* | CCATGGCTAGATCTCAGTCGACatgattatctatacata cacatacac | NST3 terminator |
| *NST3rev3* | ACTAGGTACCacggattcaccatgtgcgtt | NST3 terminator |
| *H4GFPfor4* | ACTAACATGTCTtcgggtcgtggaaaggga | Cloning H4GFP into NST3 construct, PCR for H4-GFP |
| *H4GFP-WOX4rev* | ACTAGGATCCttatttgtatagttcatccatgc | Cloning H4GFP into NST3 construct |
| *VND7for2* | ACTAGCGGCCGCtagcacgttgacgtatgctgag | VND7 promoter |
| *VND7rev2* | ACTAGGATCCatC**CATGGccacgatgat**cctataaacgt | VND7 promoter |
| *VND7for3* | ACTAGGATCCgaT**CTAGAttaaaaaaaa**acacacttctat atattg | VND7 terminator |
| *VND7rev3* | ACTAGGTACCccttctacgtggatcagaagg | VND7 terminator |
| *H4GFPfor5* | **ATCATCGTGGCCATG**tcgggtcgtggaaaggga | Cloning H4GFP into VND7 construct |
| *H4GFPrev5* | **TTTTTTTTAATCTAG**Agttataaccagtattatttgtatagttc atccatgc | Cloning H4GFP into VND7 construct |
| *H4GFP-APLfor* | ACTAACatgtcgggtcgtggaaaggga | Cloning H4GFP into APL construct |
| *H4GFP-APLrev* | ACTACTGCAGttatttgtatagttcatccatgc | Cloning H4GFP into APL construct |
| *SCRprom1* | ACTAGCGGCCGCcgaccaccaccgtcaacaat | SCR promoter |
| *SCR_Prom_R* | ACTAGGATCCAGATGcatggagattgaagggttgttggtc | SCR promoter |
| *SCR_Prom3´_F* | ACTACCCGGGcagcttggacgcctcgttcttag | SCR terminator |
| *SCR_Prom3´_R* | AGAAATGAATTCagagctccacggtggtgg | SCR terminator |
| *H4GFP-SCRfor* | ACTAGGATCCttcgggtcgtggaaaggga | Cloning H4GFP into SCR, LTP1 construct |
| *H4GFP-SCRrev* | ACTACCCGGGttatttgtatagttcatccatgc | Cloning H4GFP into SCR, LTP1 construct |
| *LTP1pGreenfor* | ACTAGCGGCCGCgaccaaaatgattaac | LTP1 promoter |
| *LTP1pGreenrev* | ACTAGGATCCatccatggattgatctcttaggtagt | LTP1 promoter |
| *LTP1prom3'for* | ACTACCCGGGtgagctagcaacggtgagatgatg | LTP1 terminator |
| *LTP1prom3'rev* | ACTAGGTACCcttttgtggaaatagaaacttggagtta | LTP1 terminator |
| *GFPrev3* | tcctctcctgcacgtatccc | PCR for H4-GFP |
| *AT5G20250pro-F* | AACAGGTCTCAACCT ttcgtgtaggcgtgtagcactcg | AT5G20250 promoter |
| *AT5G20250pro-R* | AACAGGTCTCATGTT ttttgtttctctctctctctctttggctttc | AT5G20250 promoter |
| *AT1G29520pro-F* | AACAGGTCTCAACCT attttctctgttagtagttagttaattacctaatcttagtctc | AT1G29520 promoter |
| *AT1G29520pro-R* | AACAGGTCTCATGTT ttcgccttttgaaaaactacgaaagcg | AT1G29520 promoter |
| *AT5G28630pro-F* | AACAGGTCTCAACCT aattaagccttatgtttggagcaatgtaaaaggtag | AT5G28630 promoter |
| *AT5G28630pro-R* | AACAGGTCTCATGTT cctctttctctttttttatttcttggtgtttttagttc | AT5G28630 promoter |
| *AT5G15970pro-F* | AACA GGTCTC A ACCT aaaacgcaaagaaaactttatacagtacatactcttatg | AT5G15970 promoter |
| *AT5G15970pro-R* | AACA GGTCTC A TGTT cagatatttttttcttgtaaaatcgtttttgatgtgtgttttg | AT5G15970 promoter |
| *AT2G26150pro-F* | AACA GGTCTC A ACCT tcgttagaaatgggcttaagtaaaggccc | AT2G26150 promoter |
| *AT2G26150pro-R* | AACA GGTCTC A TGTT tttcgttgtttatctcaaatccataagctcagag | AT2G26150 promoter |
| *AT1G12200pro-F* | AACA GGTCTC A ACCT | AT1G12200 promoter |

| | | |
|---|---|---|
| | taaatatataatctacggttgtagtgttttgtccaaaag | |
| *AT1G12200pro-R* | AACA GGTCTC A TGTT gttaggttatagtgaagttttgtgatctaatgaagaaag | AT1G12200 promoter |
| *AT1G24575pro-F* | AACA GGTCTC A ACCT atattatacttgcgcctcaagatgttggc | AT1G24575 promoter |
| *AT1G24575pro-R* | AACA GGTCTC A TGTT tcttaaactagctctttcttgtttggatcg | AT1G24575 promoter |
| *AT5G16010pro-F* | AACA GGTCTC A ACCT tccaactccacaagttaaaaattttcattaatgc | AT5G16010 promoter |
| *AT5G16010pro-R* | AACA GGTCTC A TGTT tttcttttgtttttggtttctcttcggaagaaag | AT5G16010 promoter |
| *AT5G15870pro-F* | AACA GGTCTC A ACCT aacactcaaaatggcttttactttctagtatcc | AT5G15870 promoter |
| *AT5G15870pro-R* | AACA GGTCTC A TGTT tttgcttagtcaagaggagaagagataacg | AT5G15870 promoter |

ATGC (underline) Sequence for restriction enzyme.
**ATGC** (bold) Sequence for In-Fusion reaction.
ATGC (uppercase) Sequence added for cloning.
atgc (lowercase) Sequence for annealing.

**Supplemental Table 2. Basic statistics of RNA-seq datasets from this study**

| *Region* | *ID* | *Raw reads* | *After trim* | *Uniquely mapped* | *Multiply mapped* | *Unmapped* | *Accession Number* |
|---|---|---|---|---|---|---|---|
| *NST3* | S1 | 33,098,137 | 59.54% | 58.13% | 23.19% | 18.68% | GSM4217858 |
| *NST3* | S2 | 26,554,872 | 46.06% | 39.04% | 23.10% | 37.85% | GSM4217859 |
| *NST3* | S3 | 33,622,145 | 72.47% | 71.40% | 19.80% | 8.81% | GSM4217860 |
| *VND7* | S4 | 53,508,021 | 35.45% | 54.56% | 36.59% | 8.85% | GSM4217867 |
| *VND7* | S5 | 28,049,974 | 40.18% | 49.02% | 41.24% | 9.74% | GSM4217868 |
| *VND7* | S6 | 21,234,034 | 44.60% | 60.59% | 28.40% | 11.01% | GSM4217869 |
| *PXY* | S7 | 42,073,803 | 79.76% | 53.76% | 29.61% | 16.63% | GSM4217870 |
| *PXY* | S8 | 52,535,458 | 82.56% | 54.35% | 23.82% | 21.83% | GSM4217871 |
| *PXY* | S9 | 20,301,474 | 77.91% | 53.76% | 26.99% | 19.25% | GSM4217872 |
| *SMXL5* | S10 | 23,176,874 | 37.69% | 57.31% | 27.95% | 14.75% | GSM4217861 |
| *SMXL5* | S11 | 23,424,841 | 43.37% | 56.51% | 30.33% | 13.16% | GSM4217862 |
| *SMXL5* | S12 | 19,248,090 | 65.06% | 66.42% | 21.85% | 11.73% | GSM4217863 |
| *APL1* | S13 | 68,143,227 | 84.51% | 55.39% | 39.61% | 5.00% | GSM4217864 |
| *APL2* | S14 | 35,571,237 | 84.91% | 60.55% | 34.78% | 4.67% | GSM4217865 |
| *APL3* | S15 | 48,232,939 | 81.70% | 70.68% | 24.22% | 5.09% | GSM4217866 |
| *SCR* | S16 | 27,756,903 | 31.27% | 40.23% | 40.94% | 18.83% | GSM4217873 |
| *SCR* | S17 | 12,568,312 | 53.14% | 51.52% | 36.81% | 11.67% | GSM4217874 |
| *SCR* | S18 | 14,885,558 | 72.22% | 57.15% | 35.60% | 7.25% | GSM4217875 |
| *LTP1* | S19 | 64,699,969 | 22.89% | 33.01% | 33.66% | 33.33% | GSM4217876 |
| *LTP1* | S20 | 16,682,896 | 35.87% | 50.14% | 26.29% | 23.57% | GSM4217877 |
| *LTP1* | S21 | 35,493,342 | 28.41% | 42.67% | 25.75% | 31.58% | GSM4217878 |
| *LCMCtl* | S30 | 23,403,325 | - | 95.51% | 1.38% | 3.11% | GSM4217888 |
| *LCMCtl* | S31 | 24,831,655 | - | 95.80% | 1.25% | 2.94% | GSM4217889 |
| *Pcap* | S32 | 23,446,707 | - | 87.50% | 1.28% | 11.22% | GSM4217890 |
| *Pcap* | S33 | 23,226,571 | - | 93.83% | 1.47% | 4.70% | GSM4217891 |
| *Pith* | S34 | 24,593,314 | - | 88.65% | 1.98% | 9.37% | GSM4217892 |
| *Pith* | S35 | 23,997,414 | - | 95.35% | 1.47% | 3.18% | GSM4217893 |
| *GFP negative* | | | | | | | |
| *SMXL5* | S40 | 17,901,192 | 42.59% | 58.18% | 20.05% | 21.77% | GSM4217882 |
| *SMXL5* | S41 | 20,436,609 | 19.77% | 37.51% | 17.76% | 44.72% | GSM4217883 |
| *SMXL5* | S42 | 18,359,819 | 28.85% | 48.91% | 18.33% | 32.76% | GSM4217884 |
| *APL* | S43 | 11,087,751 | 87.91% | 59.77% | 34.34% | 5.88% | GSM4217885 |
| *APL* | S44 | 9,220,810 | 79.36% | 47.06% | 38.86% | 14.08% | GSM4217886 |
| *APL* | S45 | 16,792,527 | 82.77% | 58.36% | 35.77% | 5.86% | GSM4217887 |
| *NST3* | S46 | 24,937,918 | 55.18% | 49.67% | 26.39% | 23.94% | GSM4217879 |
| *NST3* | S47 | 15,052,250 | 44.21% | 15.27% | 5.13% | 79.60% | GSM4217880 |
| *NST3* | S48 | 34,229,346 | 72.57% | 64.71% | 23.63% | 11.66% | GSM4217881 |

### Dataset legends

**Supplemental Dataset 1. Significantly differentially expressed genes in transcriptome datasets obtained from GFP-positive comparing to GFP-negative nuclei from *NST3$_{pro}$*, *SMXL5$_{pro}$* and *APL$_{pro}$* lines, respectively.**
Output of DESeq2 using Wald test is shown. "*baseMean*" : mean value of normalized read counts. "*log2FoldChange*" : log2 value of fold change of normalized read counts in datasets from GFP-positive nuclei compared to GFP-negative nuclei. "*lfcSE*" : the standard error estimate for the log2 fold change estimate. "*stat*" : Wald statistic. "*pvalue*" : Wald test p-value. "*padj*" : Benjamini-Hochberg adjustment of *p* value.

**Supplemental Dataset 2. Raw read counts for each FANS-derived RNA-seq dataset**
Raw read counts for all Arabidopsis genes in all dataset obtained (All_dataset) and in the reduced dataset (Reduced dataset). The name of dataset corresponds to Figure 3 and Supplemental Figure 3.

**Supplemental Dataset 3. Raw and normalized read counts for each LCM-derived RNA-seq dataset.**
Raw and normalized read counts in DESeq2 using total counts in each LCM RNA-seq dataset.

**Supplemental Dataset 4. The result of LRT analysis of RNA-seq datasets obtained from *NST3$_{pro}$*, VND7$_{pro}$, *PXY$_{pro}$*, *SMXL5$_{pro}$*, *APL$_{pro}$*, *SCR$_{pro}$* and *LTP1$_{pro}$* –positive nuclei**
Output of DESeq2 using LRT is shown. Please see the legend of Supplemental Dataset 1. Here, "log2FoldChange" represents the changes between the model with the group of each line and the reduced model. Genes with padj values lower than 0.01 are selected in one table ('SDE genes') and the results of all genes are shown in a second table ('all genes') for reference.

**Supplemental Dataset 5. Normalized read counts for each FANS-derived RNA-seq dataset**
Normalized read counts in DESeq2 using total counts in each dataset. SDE genes are shown in one table ('SDE genes') and the results for all genes are shown in a second table ('all genes') for reference.

**Supplemental Dataset 6. Clustering of genes based on their expression pattern among seven tissues.**
SDE genes were clustered and the cluster ID was assigned to each gene.

**Supplemental Dataset 7. Average value of normalized read counts of SDE genes within each nuclei type, ranked by the values.**
Normalized read counts (Supplemental Dataset 5) were averaged for each tissue. The average value and the rank of each tissue are presented. Data are sorted according to the ratio of the highest value to the second highest value.

**Supplemental Dataset 8. SDE genes comparing Phloem cap/Pith and the remaining vascular bundle in LCM-derived datasets.**
Output of DESeq2 using Wald test is shown. Please see the legend of Supplemental Dataset 1. Here, "*log2FoldChange*" indicates the log2 value of the fold change of normalized read counts in phloem cap / pith datasets compared with the remaining vascular bundle. Genes with *padj* > 0.01, *log2FoldChange* >1 were selected, sorted according to log2FoldChange and shown for phloem cap and pith, respectively.

**Supplemental Dataset 9. GO term enrichment analysis in phloem cap-associated or pith-associated genes.**
Output of PANTHER Overrepresentation Test using Fisher's Exact test with Bonferroni correction is shown. For a detailed description of each column, please see the help option of PANTHER (http://go.pantherdb.org/tips/tips_overrep.jsp). „Arabidopsis thaliana - REFLIST" indicates the number of genes registered for each GO term. "phloem cap "or "pith" indicates the number of genes for each GO term in the list of interest. "phloem cap (expected)" indicates the expected number of genes for each GO term. "fold Enrichment" indicates the fold enrichment of the genes observed in the list of interest over the expected value. "over/under" indicates overrepresented or underrepresented terms in the list. Here, only overrepresented terms were selected. "P-value" : *p* value in Fisher's Exact test with Bonferroni correction corresponds to the false discovery rate. GO terms with P-value less than 0.05 are shown.

**Supplemental Dataset 10. SDE genes comparing *$NST3_{pro}$*-positive and *$VND7_{pro}$*-positive nuclei**
Output of DESeq2 using Wald test is shown. Please see the legend of Supplemental Dataset 1. Here, "*log2FoldChange*" indicates the log2 value of fold change of normalized read counts from *$VND7_{pro}$*-positive nuclei compared to *$NST3_{pro}$*-positive nuclei. 14,063 SDE genes were compared and genes with *padj* > 0.01 and absolute value of *log2FoldChange* > 1 were selected and sorted according to the log2FoldChange. Thus, genes are ordered from *$VND7_{pro}$* enriched (top) to *$NST3_{pro}$* enriched (bottom).

**Supplemental Dataset 11. GO term enrichment analysis among genes specifically active in *$NST3_{pro}$*-positive when compared to *$VND7_{pro}$*-positive nuclei, and among genes specifically active in *$VND7_{pro}$*-positive nuclei compared to *$NST3_{pro}$*-positive nuclei.**
The output of PANTHER Overrepresentation Test using Fisher's Exact test with Bonferroni correction is shown. Please see the legend of Supplemental Dataset 9.

**Supplemental Dataset 12. SDE genes comparing *$PXY_{pro}$*-positive nuclei and *$SMXL5_{pro}$*-positive nuclei.**

The output of DESeq2 using Wald test is shown. Please see the legend of Supplemental Dataset 1. Here, "*log2FoldChange*" indicates the log2 value of the fold change of normalized read counts in $SMXL5_{pro}$-positive nuclei compared to $PXY_{pro}$-positive nuclei. 14,063 SDE genes were compared and genes with *padj* > 0.01 and absolute value of *log2FoldChange* >1 were selected and sorted according to log2FoldChange. Thus, genes are sorted from $SMXL5_{pro}$ enriched (top) to $PXY_{pro}$ enriched (bottom).

**Supplemental Dataset 13. GO term enrichment analysis among genes specifically active in $PXY_{pro}$-positive when compared to $SMXL5_{pro}$-positive nuclei, and among genes specifically active in $SMXL5_{pro}$-positive when compared to $PXY_{pro}$-positive nuclei.**
The output of PANTHER Overrepresentation Tests using Fisher's Exact test with Bonferroni correction is shown. Please see legend of Supplemental Dataset 9.

**Supplemental Dataset 14. Fold enrichment values of significantly enriched transcription factor binding regions in the upstream regions of genes from FANS/RNA-seq-derived clusters and tissue-specific genes from LCM/RNA-seq analyses.**
Shown are fold enrichment values of each transcription factor binding region in FANS/RNA-Seq clusters and in LCM-derived phloem cap and pith-associated genes. AGI_TF indicates the Arabidopsis Genome Initiative (AGI) codes for each associated transcription factor. DAP_ID indicates the binding region ID determined by DNA affinity purification sequencing (O'Malley et al., 2016). Only significantly enriched regions are shown (p < 8.8e-05 (Bonferroni adjusted threshold of 0.05) in Fisher's exact test).
